# Macropinocytosis inhibition attenuates pro-fibrotic responses in lung fibroblasts and pulmonary fibrosis

**DOI:** 10.1101/2025.07.09.663937

**Authors:** Ivan O. Rosas, Aaron K. McDowell-Sanchez, Santiago Sanchez, Juan D. Cala-Garcia, Alan R. Waich Cohen, Maria Elisa Ruiz Echartea, Scott A. Ochsner, Daniel Kraushaar, Lindsay J. Celada, Dandan Sun, Francesca Polverino, Cristian Coarfa, Neil J. McKenna, Konstantin Tsoyi

## Abstract

Idiopathic pulmonary fibrosis (IPF) is a devastating chronic lung disorder with limited treatment options. Macropinocytosis is one of the key cellular processes involved in nutrient consumption from the extracellular environment under stress conditions. Here, we studied the role of macropinocytosis in lung fibroblast activation and experimental pulmonary fibrosis. We found that macropinocytosis is increased in human lung fibroblasts (HLFs) derived from IPF patients. The inhibition of macropinocytosis with 5-(n-ethyl-n-isopropyl)-amiloride (EIPA) significantly inhibited profibrotic responses in IPF-derived and TGF-β1-stimulated HLFs. EIPA exerted antifibrotic effects by regulating amino acid (AA) uptake, mammalian target of rapamycin complex 1 (mTORC1) activation and mesenchyme homeobox1 (MEOX1) expression in activated HLFs. Both genetic and pharmacological inhibition of macropinocytosis significantly ameliorated pulmonary fibrosis in bleomycin (Bleo)-injured mice. Using IPF-derived precision cut lung slices (PCLS), we observed robust repression of profibrotic gene expression programs in EIPA-treated PCLS across different fibroblast subpopulations. Finally, we found that imipramine (Imi), a tricyclic antidepressant approved by the Food and Drug Administration (FDA), effectively inhibited macropinocytosis and ameliorated profibrotic responses in lung fibroblasts, Bleo-injured mice and IPF-derived PCLS. Taken together, our results suggest macropinocytosis inhibition as a potential therapeutic strategy to treat pulmonary fibrosis.

## INTRODUCTION

Idiopathic pulmonary fibrosis (IPF) is a chronic, progressive, fibroproliferative interstitial lung disease of enigmatic etiology (1). IPF affects approximately 500 per 100,000 adults over the age of 65 in the United States alone and is increasing in prevalence, leading to rising rates of hospital admissions and deaths (2). Currently, there are two Food and Drug Administration (FDA) approved therapies to treat IPF, nintedanib and pirfenidone, both of which can reduce decline in lung function, decrease number of acute exacerbations and improve survival (3, 4). However, neither of these drugs can stop or reverse disease progression. Thus, there is an unmet need to determine the molecular mechanisms underlying the pathogenesis of IPF to enable the development of new and effective therapeutics.

Fibroblasts/myofibroblasts are among the primary cell types responsible for the accumulation of extracellular matrix (ECM) and organ remodeling in fibrotic disorders (5). Fibroblasts can be activated by pathological stimuli such as transforming growth factor-beta (TGF-β) and substrate stiffness, prompting these cells to express increased α-smooth muscle actin (α-SMA) stress fibers, deposit ECM proteins, display increased contractility and invasiveness, and develop resistance to pro-apoptotic stimuli (*e.g.,* Fas ligand) (6, 7). Thus, targeting these processes in fibroblasts represents an opportunity for therapeutic intervention in fibrotic disorders.

Macropinocytosis is an actin-dependent, but clathrin-independent, endocytic process that mediates the nonselective internalization of extracellular materials such as proteins, cell debris, or viruses (8). Macropinosomes are formed when cell membrane ruffles curve to form circular cup-shaped structures, which eventually seal and pinch off to form large vesicles (greater than 0.2 μm in diameter) (8). Macropinocytosis can function as a means of nutrient acquisition in mammalian cells which in turn can greatly influence cell survival and proliferation (9, 10).

Multiple reports also highlight the key role for macropinocytosis in various malignant disorders by providing essential nutrients to support the proliferation and metastasis of tumor cells under nutrient poor conditions (8, 11). However, the role of macropinocytosis in organ fibrosis remains unclear.

Here, we report that macropinocytosis inhibition exerts potent antifibrotic effects in activated lung fibroblasts by regulating amino acid (AA) uptake and mTORC1 activation. We also found that imipramine, a clinically-used tricyclic antidepressant, inhibits macropinocytosis in IPF-derived lung fibroblasts and ameliorates experimental pulmonary fibrosis in *ex vivo* and *in vivo* models.

## RESULTS

### Macropinocytosis inhibition attenuates profibrotic responses in activated HLFs

Lung fibroblasts derived from IPF lung tissue possess a higher capacity for myofibroblast differentiation, ECM production, resistance to cell death and cell invasion (12–14). We investigated whether macropinocytosis levels are elevated in IPF-derived lung fibroblasts compared to control fibroblasts. The uptake of fluorescent labeled large particles, such as 70 kDa dextran, is a widely accepted method to measure macropinocytosis in non-phagocytic cells (9, 15–17). Using fluorescence microscopy and flow cytometry techniques, IPF-derived lung fibroblasts showed increased levels of macropinocytosis compared to control cells, as measured by the uptake of 70kDa FITC-labeled dextran (**Figure 1A and B**). Unlike other types of endocytosis, macropinocytosis is dependent on Na^+^/H^+^ proton exchangers (NHEs), also known as the solute carrier 9 family (SLC9s) (18–20). Thus, 5-(N-Ethyl-N-isopropyl) amiloride (EIPA), a known pan-NHE blocker, is considered as a first choice pharmacological inhibitor of macropinocytosis in biomedical research (16, 21–25). Consistent with prior reports, EIPA effectively inhibited macropinocytosis in IPF-derived lung fibroblasts, while neither affecting clathrin-dependent endocytosis nor expression of caveolin-1, a key protein in caveolin-mediated endocytosis (**Supplemental Figure 1 A-C**). We tested whether EIPA can inhibit profibrotic responses in IPF-derived lung fibroblasts. As shown in **Figure 1 C and D**, EIPA potently downregulated collagen 1 (ECM protein) and alpha-smooth muscle actin (α-SMA), a marker of myofibroblast transformation, protein and mRNA expression in IPF-derived lung fibroblasts.

**Figure 1:**
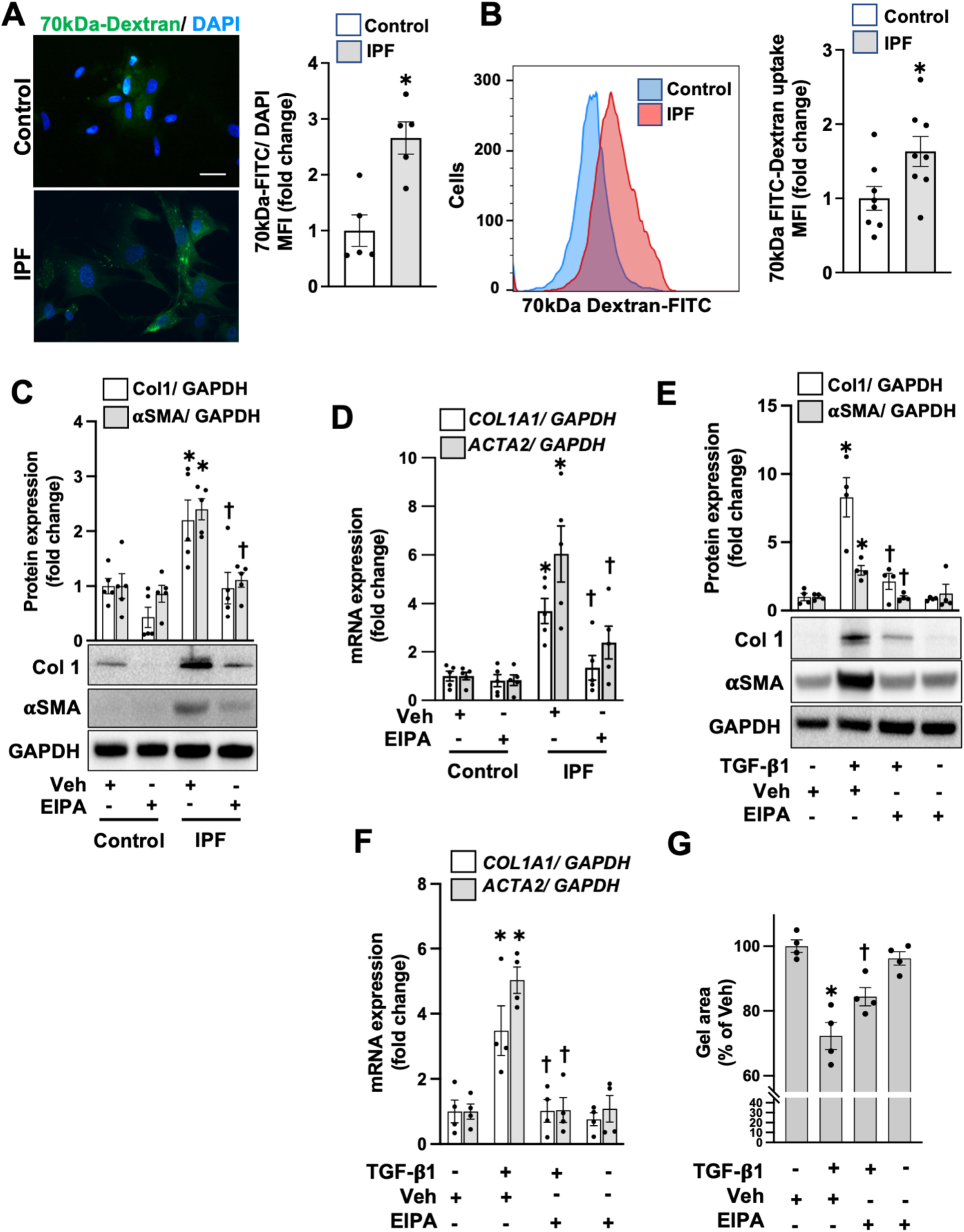
M**a**cropinocytosis **is elevated in IPF-derived lung fibroblasts and its inhibition regulates profibrotic responses.** (A and B) Control and IPF-derived lung fibroblasts were conditioned in low serum media for 24 hours as described in Methods. 24 hours later, culture media was changed and cells we incubated with FITC-Dextran (70 kDa, 0.5 mg/ml) for 1 hour. Then, cells were washed three times with PBS. Macropinocytosis was measured by fluorescent microscopy (A, scale bar represents 25 μm) and flow cytometry (B) to measure fluorescence intensity of FITC-Dextran in cells (n=5 for A, and n=8 for B in each condition). (C and D) Cells were treated with Vehicle (Veh) or EIPA (12.5 μM) for 24 hours. Then cells were harvested and subjected to western blot (C) and qRT-PCR (D) to measure collagen 1 and αSMA expression as described in Methods (n= 5 each condition). (E and F) Control HLFs were treated with EIPA (12.5 μM) with or without TGF-β1 (10 ng/ml) for 24 hours. Then cells were harvested and subjected to western blot (E) and qRT-PCR (F) as described in Methods (n=4 each condition). (G) Control HLFs were mixed with collagen 1 solution as described in Methods and treated with EIPA (12.5 μM) with or without TGF-β1 (10 ng/ml). Gel size was measured at 0 and 24 h after collagen gelation (n=4 for each condition). Data are mean ± SEM. *P* < 0.05; significant comparisons by Student t-test or one-way ANOVA: *vs. unstimulated or Control+Veh, ^†^ vs. IPF+Veh or TGF-β1 alone.

Similar results were found in TGF-β1 stimulated control lung fibroblasts, where EIPA attenuated collagen 1 production, myofibroblast transformation and cell contractility (**Figure 1 E, F and G**). Of note, we observed a mild, but not statistically significant, decrease in fibronectin 1 (*FN1*) expression in response to EIPA in activated HLFs (**Supplemental Figure 1 D**). These data suggest that macropinocytosis activity is elevated in activated or IPF-derived lung fibroblasts, and that inhibition of this process leads to decreased profibrotic responses in these cells.

### Macropinocytosis regulates profibrotic responses in activated HLFs by regulating AA-mediated mTORC1 activation

Previous reports suggest that macropinocytosis tightly regulates mTORC1 activation by regulating cytosolic AA levels, which can be obtained by direct uptake of free AAs or proteins (*e.g.,* serum albumin) from the extracellular environment (9–11, 19). In turn, mTORC1 activation is one of the central signaling pathways in promoting profibrotic responses in fibroblasts by activating multiple downstream signaling pathways and regulating cell metabolism (26–28).

Accordingly, we found that EIPA inhibits the phosphorylation of p70S6K and S6, which are representative of mTORC1 activation and metabolic reprogramming, in TGF-β1 activated lung fibroblasts in low serum (1 %) culture medium without affecting cell viability (**Figure 2 A, Supplementary Figure 2 A and B**). Interestingly, withdrawal of AAs, using AA-free cell culture media, completely abrogated mTORC1 activation even in the presence of TGF-β1 (**Figure 2 B**). The addition of AAs resulted in the upregulation of phospho-S6 in lung fibroblasts, which was further enhanced by TGF-β1 treatment (**Figure 2 B**). EIPA potently attenuated S6 activation under AA supplemented conditions (**Figure 2 B**). Further, we found that macropinocytosis regulates the amount of cytosolic AAs under these conditions, suggesting that under profibrotic conditions (e.g., under TGF-β1 stimulation), extracellular AAs play a significant role in mTORC1 activation and that macropinocytosis inhibition can significantly inhibit this process (**Figure 2 C**). Finally, we found that mTORC1 activation by TSC1 silencing, a negative regulator of mTOR, attenuated the antifibrotic effects of EIPA (**Figure 2 D**). These data suggest that macropinocytosis regulates the uptake of extracellular AAs which in turn leads to mTORC1 activation and increased profibrotic responses in lung fibroblasts.

**Figure 2:**
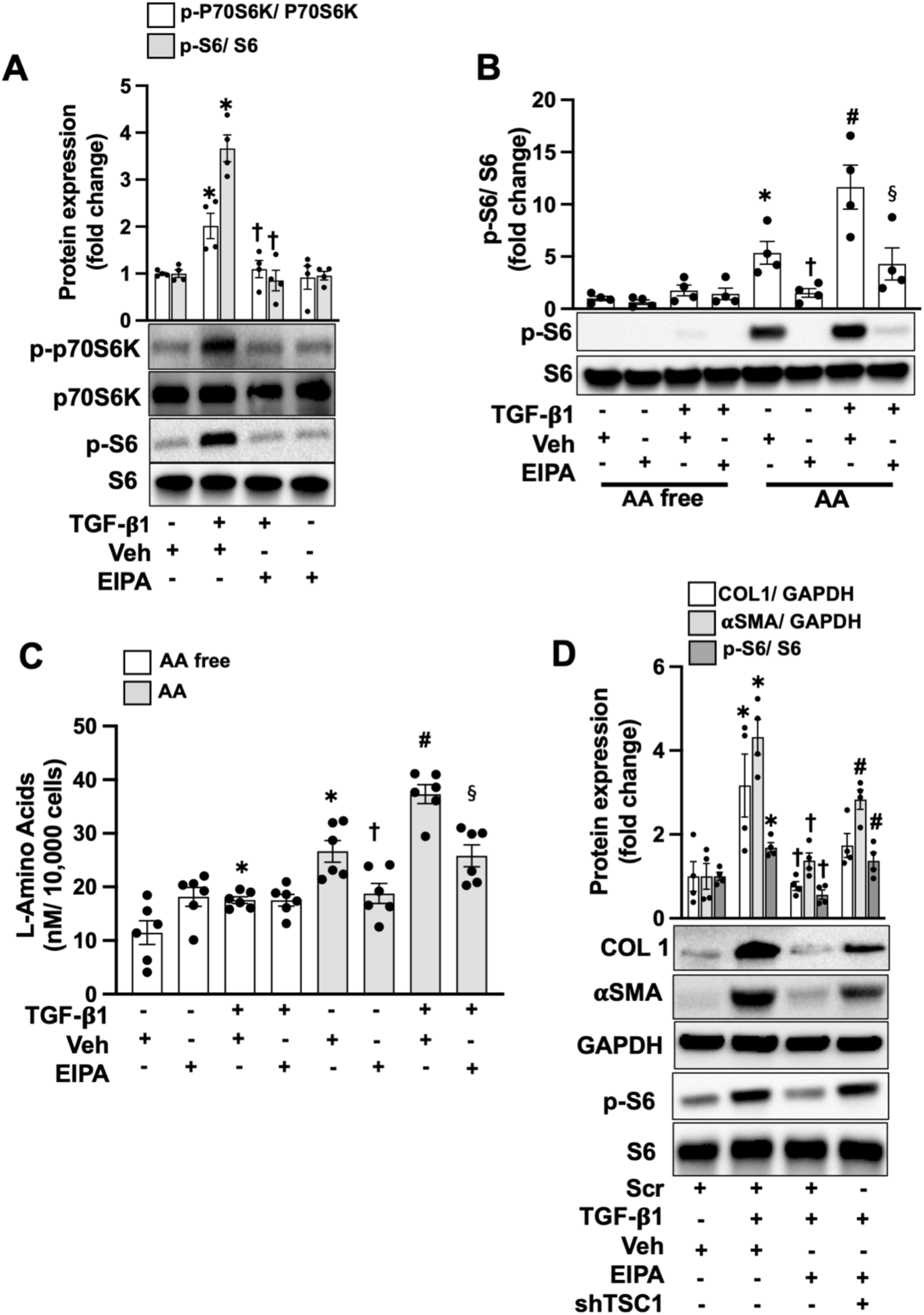
M**a**cropinocytosis **inhibition ameliorates mTORC1 signaling by regulating amino acid uptake in lung fibroblasts.** (A) Control HLFs were treated with EIPA (12.5 μM) with or without TGF-β1 (10 ng/ml) for 24 hours. Then, p-P70S6K and p-S6 were measured by western blot as described in Methods (n=4 each condition). (B and C) Cells were treated with EIPA (12.5 μM) with or without TGF-β1 (10 ng/ml) for 4 hours in AA-free and AA-supplemented conditions as described in Methods. Then, cells were harvested to measure p-S6 by western blot (B) and L-amino acid assay as described in Methods (n=4 or 6). (D) Cells were lentivirally transfected with scramble (Scr) or shTSC1 as described in Methods. Then, cells were treated with EIPA (12.5 μM) with or without TGF-β1 (10 ng/ml) for 24 hours. Then, cells were harvested and COL1, αSMA and p-S6 were measured by western blot (n=4 each condition). Data are mean ± SEM. *P* < 0.05; significant comparisons by Student t-test or one-way ANOVA: * vs. Veh or Veh + AA free, ^†^ vs. TGF-β1 + Veh or Veh + AA, ^§^ vs. TGF-β1 + Veh + AA, ^#^ vs. Scr + TGF-β1 + EIPA.

### MEOX1 is regulated by macropinocytosis and mTORC1 in activated HLFs

We next aimed to evaluate the effect of EIPA treatment on the TGFB-induced transcriptional program in MRC5 human lung fibroblasts (HLFs). Consistent with a robust repressive effect of EIPA on the TGF-β1 (TGFB)-induced transcriptional prorgram, TGFB-induced genes were strongly enriched among EIPA-repressed genes (**Figure 3 A**). TGFB-induced genes repressed by EIPA treatment included several with well established roles in lung fibrosis such as *ACTA2* and *COL1A1*. Notably, *MEOX1,* recently identified as a key transcription factor in fibroblast activation and cardiac fibrosis, was among the repressed genes (29). Accordingly, we next investigated whether MEOX1 was regulated by macropinocytosis-mediated mTORC1 activation and contributed to the profibrotic phenotype of activated HLFs. We found that both EIPA and Rapalink-1, a known mTORC1 inhibitor, downregulated MEOX1 mRNA and protein expression levels (**Figure 3 B and C**). Furthermore, silencing of regulatory-associated protein of mTOR (RAPTOR), a key component of mTORC1, but not mTORC2, inhibited MEOX1 expression in TGF-β1-activated HLFs (**Figure 3 D**). Of note, we also found that activation of mTOR by shTSC1 significantly increased MEOX1 gene expression (**Supplemental Figure 2 C**). Finally, silencing of MEOX1 potently inhibited COL1 and α-SMA expression in activated HLFs (**Figure 3 E**). These data suggest that macropinocytosis inhibition negatively affects profibrotic responses in fibroblasts by regulating an mTORC1/ MEOX1 signaling axis.

**Figure 3.**
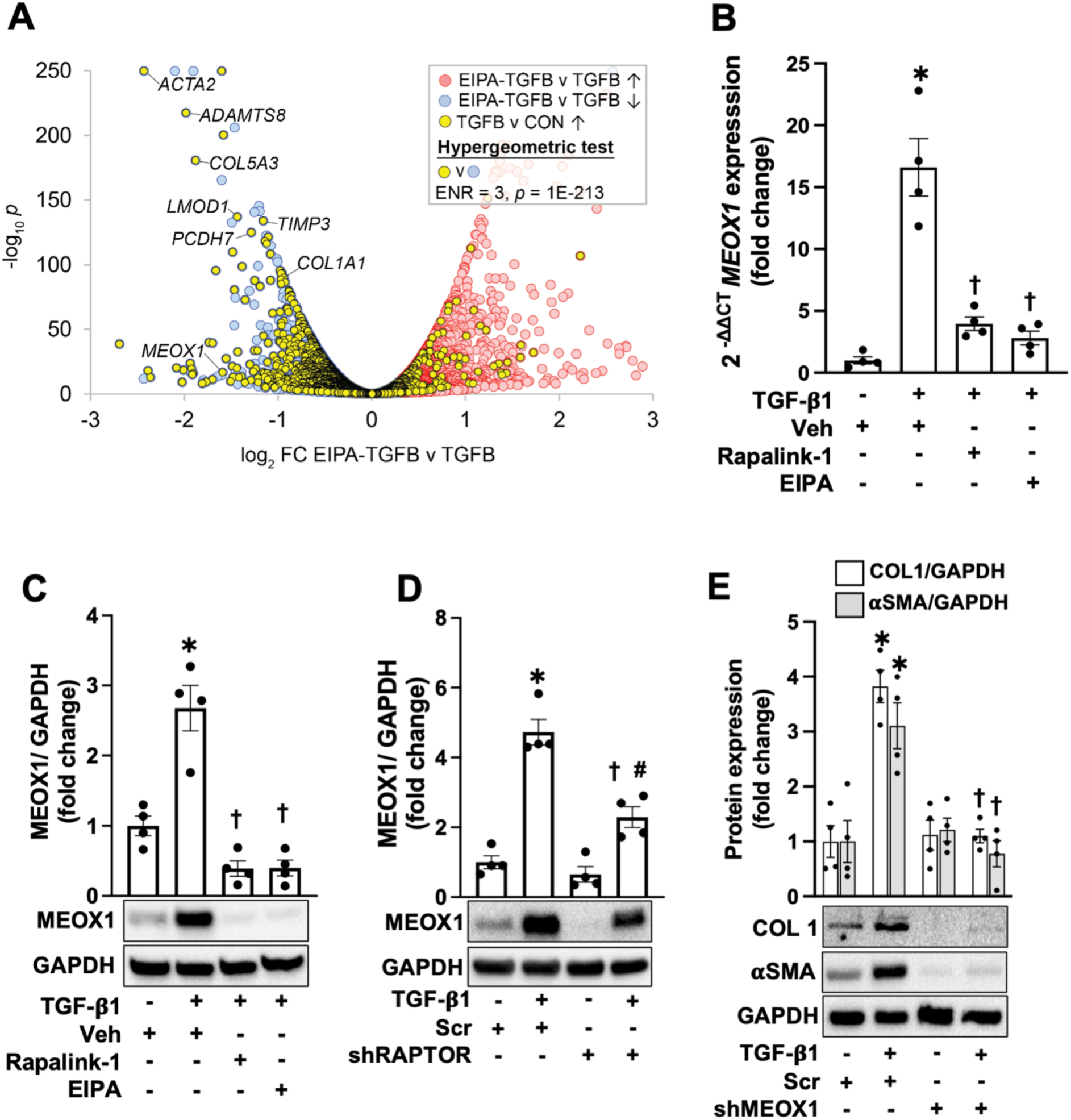
Macropinocytosis inhibition regulates MEOX1 expression in activated lung fibroblasts. (A) Volcano plot representing treatment of MRC5 HLFs with and without EIPA in the presence of TGFB. Overlaid in yellow is a set of TGFβ1-induced genes from a separate RNA-Seq experiment in MRC5 HLFs treated with TGF-β1 (10 ng/ml) or vehicle. MRC-5 cells were treated with EIPA (12.5 μM) with or without TGF-β1 (10 ng/ml) for 24 hours. Then, RNA was isolated and subjected to bulk RNA sequencing as described in Methods (n=3). (B and C) Control HLFs were treated with EIPA (12.5 μM), and Rapalink-1 (10 nM) with or without TGF-β1 (10 ng/ml) for 24 hours. Then, MEOX1 at mRNA and protein levels were measured by qRT-PCR (B) and western blot (C) respectively (n=4 each condition). (D) Control HLFs were lentivirally transfected with Scr or shRAPTOR as described in Methods. Then, cells were treated with TGF-β1 (10 ng/ml) for 24 hours or left untreated. Protein levels of MEOX1 were measured by western blot as described in Methods (n=4 each condition). (E) Control HLFs were lentivirally transfected with Scr or shMEOX1 as described in Methods. Then, cells were treated with TGF-β1 (10 ng/ml) for 24 hours or left untreated. Protein levels of COL1 and αSMA were measured by western blot as described in Methods (n=4 each condition). Data are mean ± SEM. *P* < 0.05; significant comparisons by Student t-test or one-way ANOVA: * vs. unstimulated or Scr alone, ^†^ vs. TGF-β1 alone.

### Macropinocytosis inhibition attenuates lung fibrosis in Bleo-injured mice

Next, we examined whether genetic silencing or pharmacological inhibition of macropinocytosis can ameliorate pulmonary fibrosis in Bleo-injured mice. Previously, Nhe1/Slc9a1 deficiency was shown to markedly inhibit macropinocytosis *in vivo* (30). Indeed, we found that Slc9a1 deficient lung fibroblasts exhibited decreased macropinocytosis activity compared to their wild type (WT) counterparts, without affecting clathrin-dependent endocytosis or caveolin1 expression (**Supplemental Figure 3**). Next, we found that deficiency of Slc9a1 in fibroblasts significantly attenuated pulmonary fibrosis in mice subjected to intratracheal Bleo administration (**Figure 4 A and B, Supplemental Figure 4 A**). Consistently, delayed administration of a macropinocytosis inhibitor (EIPA, 10 mg/kg) significantly reduced lung fibrosis as determined by Trichrome and H&E staining and hydroxyproline assay, without affecting lung inflammation in Bleo-injured mice (**Figure 4 C and D, Supplemental Figure 4 B, 5 B and C**). Accordingly, total lung homogenates showed a significant downregulation of profibrotic gene expression, and MEOX1 expression in fibroblast-specific Slc9A1 deficient and EIPA-treated mice (**Figure 4 E and F**).

**Figure 4.**
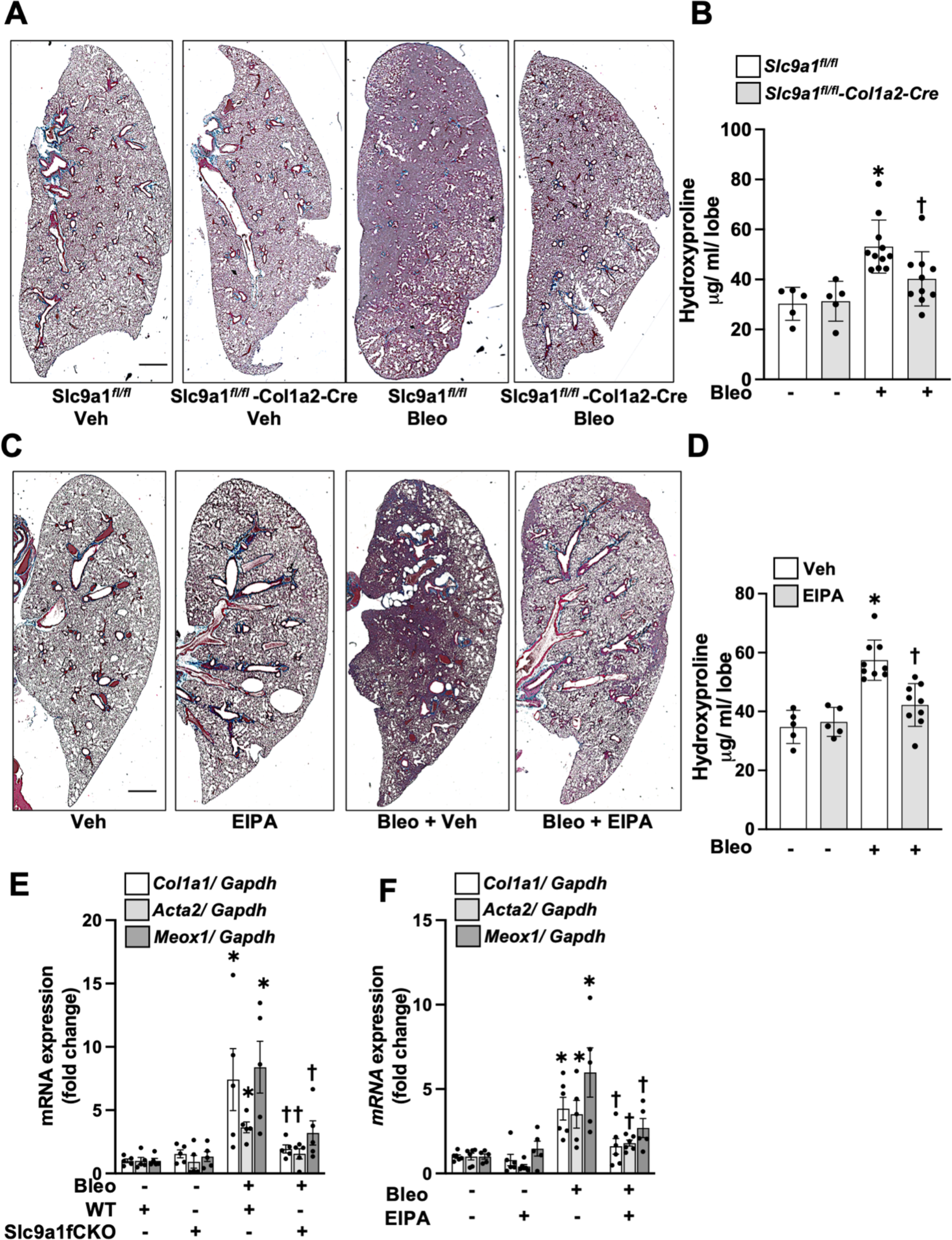
Macropinocytosis inhibition attenuates pulmonary fibrosis in Bleo-injured mice. (A and B) *Slc9a1^fl/fl^* (WT) or *Slc9a1^fl/fl^* Col1a2^Cre-ER(T)+/0^ (Slc9a1 fCKO) were treated with tamoxifen to induce Slc9a1 knockout in fibroblasts as described in Methods. Then, mice were exposed to Bleo-induced pulmonary fibrosis as described in Methods. At day 21 after Bleo exposure, lungs were harvested, stained with Masson’s trichrome staining (A) scale bar represents 1000 μm, and subjected to hyproxyproline assay (B) (n=5 for sham groups, n=10-11 for Bleo groups). (C and D) WT mice were exposed to Bleo to induce pulmonary fibrosis as described in Methods. 10 days after Bleo exposure, mice were intraperitoneally treated with EIPA (10 mg/kg) or vehicle (Veh) every other day until the end of experiment. At day 21 after Bleo exposure, lungs were harvested, stained with Masson’s trichrome staining (n=3-4) (C), and subjected to hyproxyproline assay (D) (n=5 for sham groups, n=9 for Bleo groups). E) The lungs from WT and Slc9a1 fCKO mice were subjected to RNA isolation as described in Methods. The mRNA levels of Col1a1, Acta2 and Meox1 were measured with qRT-PCR as described in Methods (n=5 each condition). (F) The lungs from WT mice exposed to EIPA treatment with or without Bleo injury were subjected to RNA isolation as described in Methods. The mRNA levels of Col1a1, Acta2 and Meox1 were measured with qRT-PCR as described in Methods (n=5 each condition). Data are mean ± SEM. *P* < 0.05; significant comparisons by one-way ANOVA: *vs. No Bleo (Sham), ^†^vs. Bleo alone.

Taken together, our data strongly suggest that macropinocytosis inhibition ameliorates pulmonary fibrosis in mice.

### Macropinocytosis inhibition regulates the expression of profibrotic genes in different fibroblast subpopulations in IPF-derived PCLS

Pulmonary fibroblasts represent a heterogenous cell population. To determine the effect of macropinocytosis inhibition on different fibroblast subpopulations in IPF lung, we treated PCLS derived from IPF subjects with EIPA and compared them with vehicle-treated PCLS from the same subjects. Single cell RNA sequencing (sc-RNA-seq) identified seven cell populations of mesenchyme origin (**Figure 5 A, Supplemental Figure 6**). Consistent with the results obtained from isolated fibroblasts and Bleo-injured mice, we observed a potent repressive effect of EIPA on the IPF-induced transcriptional program in alveolar fibroblasts (**Figure 5 B),** recently found to contribute to the progression of pulmonary fibrosis in mice and IPF patients (31).

**Figure 5.**
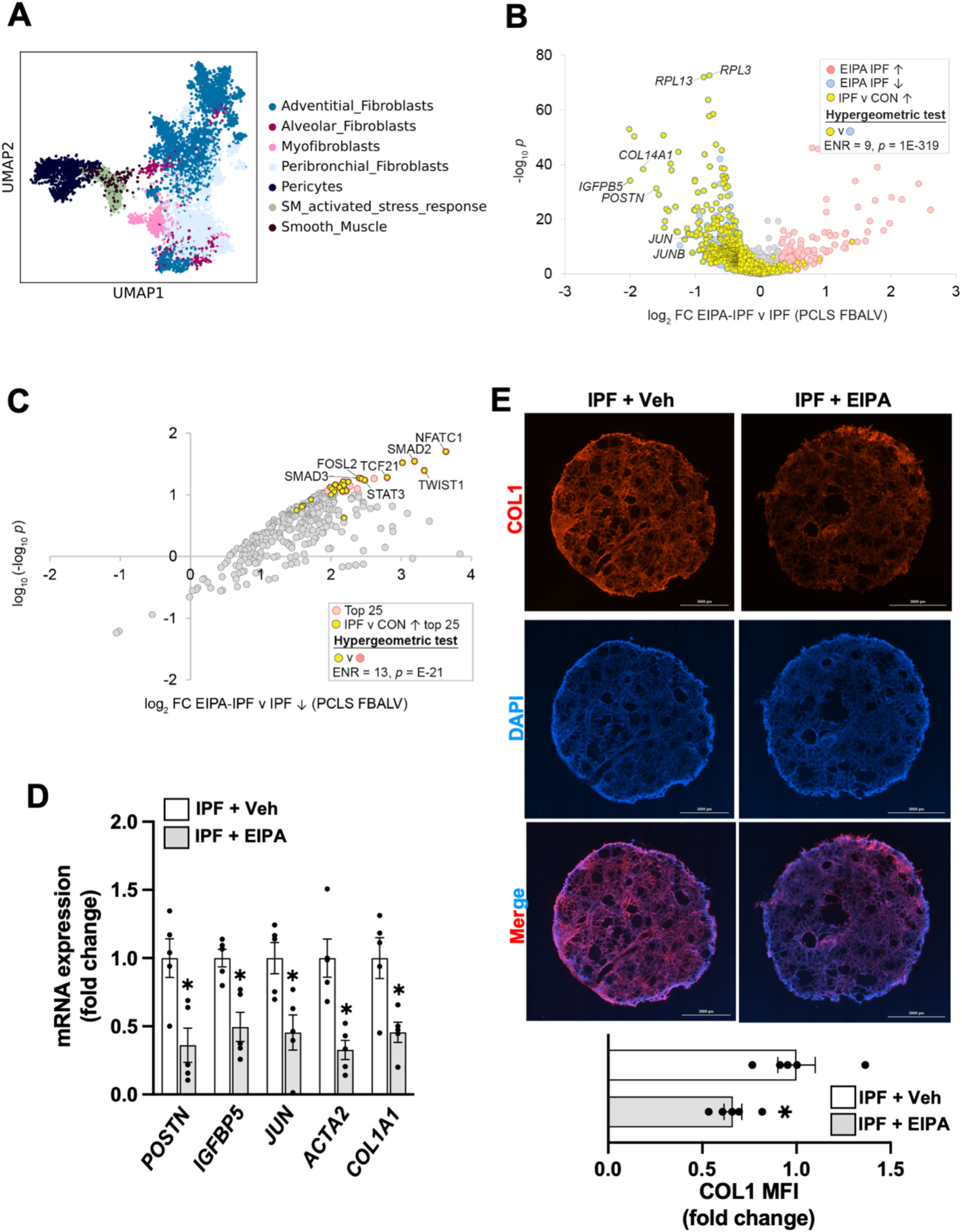
EIPA inhibits profibrotic responses in IPF-derived PCLS. IPF-derived PCLS were treated with EIPA (12.5 μM) as described in Methods. A) UMAP plot represents different mesenchymal cell populations identified in IPF PCLS. B) Volcano plot representing treatment of IPF PCLS with and without EIPA. Overlaid in yellow is a set of FBALV IPF-induced genes from a separate PCLS experiment comparing IPF and control PCLS samples. (C) Regulatory network plot for genes repressed in EIPA-treated IPF PCLS compared to IPF alone. TFs with the largest and most significant footprints within the gene set are distributed towards the top right of the plot. (D) After treatment PCLS were lysed and subjected to total RNA isolation. mRNA levels of POSTN, IGFBP5, JUN, ACTA2, and COL1A1 were measured with qRT-PCR as described in Methods. (E) After treatment, PCLS were washed with PBS and fixed for COL1 staining as described in Methods. The COL1 fluorescent intensity was measured with ImageJ software (n=5 for each condition). Scale bar represents 3000 μm. Data are mean ± SEM. *P* < 0.05; significant comparisons by one-way ANOVA: *vs. No Bleo (Sham), ^†^vs. Bleo alone.

Additionally, EIPA exerted a measurable effect on the other six mesenchymal subpopulations (**Supplemental Figure 6**). To identify candidate TFs impacted by EIPA in IPF, we carried out regulatory network analysis on genes repressed by EIPA treatment of IPF PCLS samples. Having established the impact of EIPA on IPF-induced gene expression in alveolar fibroblasts, we next wished to identify transcription factors whose function was impacted by EIPA treatment in these cells. Regulatory network analysis predicts functional footprints for transcription factors within clinical gene sets of interest (32, 33). Reflecting the strong antfibrotic effect of EIPA observed at the transcriptional level, EIPA-repressed genes contained strong footprints for numerous TFs with established roles in pulmonary fibrosis, including NFATC1, TWIST1, STAT3 and members of the SMAD family (**Figure 5 C**), suggesting that EIPA may exert its antifibrotic effects by targeting one, or all, of these factors.We validated EIPA-mediated downregulation of profibrotic genes such as periostin (POSTN), insulin growth factor binding protein-5 (IGFBP5), COL1A1 and ACTA2 by RT-qPCR in PCLS homogenates (**Figure 5 D**). Finally, collagen 1 expression was also downregulated in EIPA-treated PCLS compared to vehicle-treated PCLS (**Figure 5 E**). **Imipramine inhibits fibroblast activation and pulmonary fibrosis.**

A previous report suggested that imipramine (Imi) selectively inhibits macropinocytosis over other types of endocytosis in macrophages (21). We also found that imipramine strongly inhibited macropinocytosis in IPF-derived lung fibroblasts without affecting clathrin-dependent endocytosis or caveolin-1 expression (**Supplemental Figure 1A**). Imipramine effectively inhibited profibrotic responses in IPF-derived and TGF-β1-induced lung fibroblasts (**Figure 6 A-C**). Similar to EIPA, imipramine significantly ameliorated pulmonary fibrosis in Bleo-injured mice and profibrotic responses in IPF-derived PCLS (**Figure 5 D-G, Supplemental Figure 5**).

**Figure 6.**
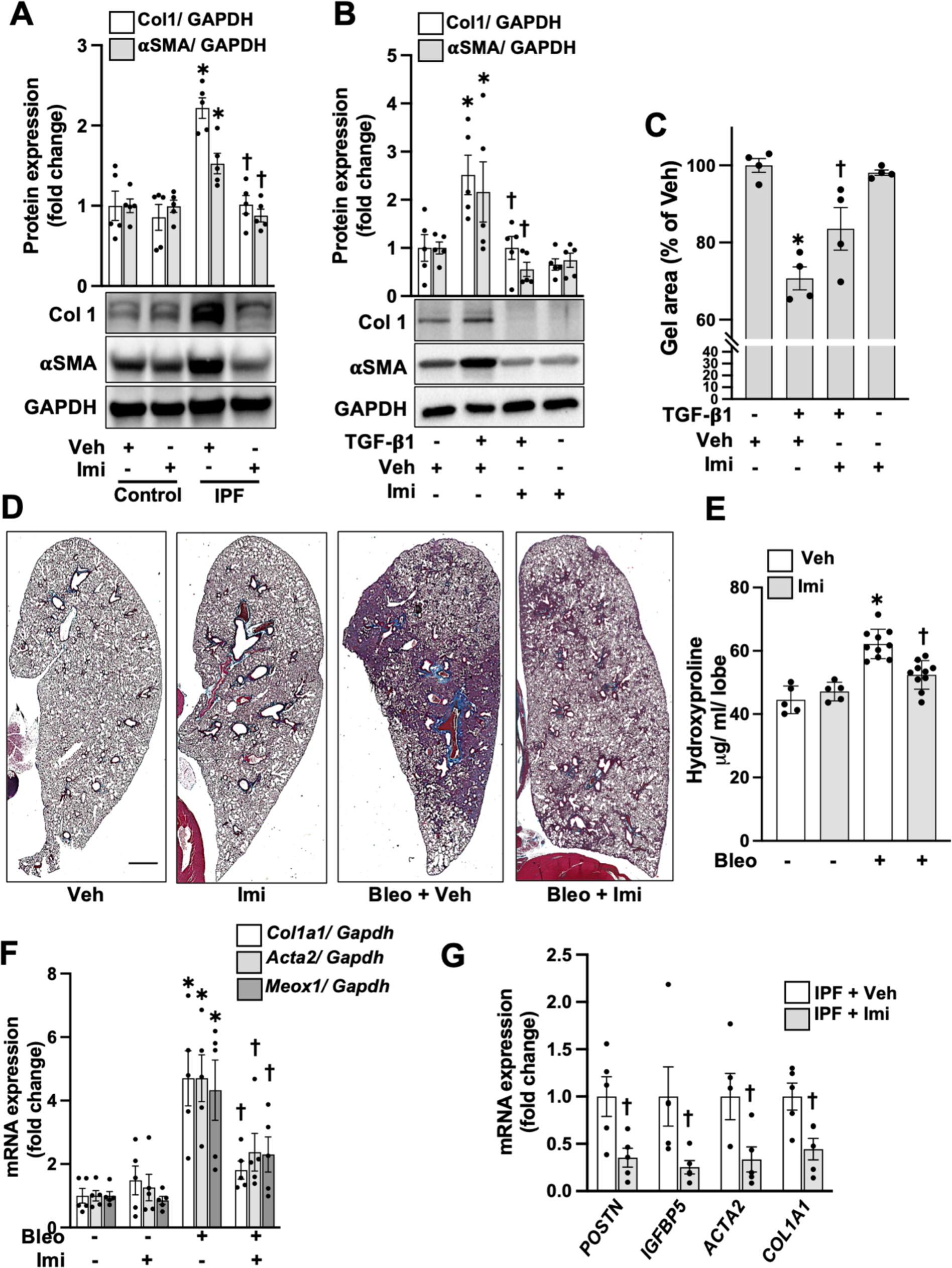
Imipramine ameliorates profibrotic responses in lung fibroblasts, Bleo-injured mice and PCLS. (A) Control and IPF derived lung fibroblasts were treated with imipramine (Imi, 10 μM). 24 hours later cells were harvested and COL1 and αSMA were measured with western blot (n=4 each condition). (B) Control HLFs were treated with Imi (10 μM) with or without TGF-β1 (10 ng/ml) for 24 hours. Then cells were harvested and subjected to western blot to measure COL1 and αSMA (n=4 for each condition). (C) Control HLFs were mixed with collagen 1 solution as described in Methods and treated with Imi (10 μM) with or without TGF-β1 (10 ng/ml). Gel size was measured at 0 and 24 h after collagen gelation (n=4 for each condition). (D, E and F) WT mice were exposed to Bleo-induced pulmonary fibrosis as described in Methods. 10 days after Bleo exposure mice were intraperitoneally treated with Imi (10 mg/kg) or vehicle (Veh) every other day until the end of experiment. At day 21 after Bleo exposure, lungs were harvested, stained with Masson’s trichrome staining (n=3-4) (D), and subjected to hyproxyproline assay (n=5 or 10) (E) or mRNA assessment (n=5) (F). (G) IPF-derived PCLS were treated with Imi (10 μM) as described in Methods. Then, cells were lysed and mRNA levels of POSTN, IGFBP5, COL1A1, and ACTA2 were measured by qRT-PCR. *P* < 0.05; significant comparisons by one-way ANOVA: * vs. unstimulated or Control or No Bleo (Sham), ^†^ vs. TGF-β1 or IPF alone or Bleo alone.

Together, these results suggest that antifibrotic effects in IPF can be achieved by targeting macropinocytosis either with EIPA or through the potential repurposing of the FDA-approved drug imipramine.

## DISCUSSION

The observation that fibroblasts may undergo membrane ruffling followed by endocytosis has been documented as early as the mid-1980s (34). Furthermore, these studies have suggested that macropinocytosis in fibroblasts can be activated by either genetic intervention (*e.g.,* Ras and Src pathway activation), or by growth factors such as platelet-derived growth factor (PDGF) (34–36). More recently, Zhang *et al.* demonstrated that TGF-β1, a potent profibrotic growth factor, induces macropinocytosis in pancreatic ductal adenocarcinoma (PDAC) cells (37). Despite multiple reports suggesting that macropinocytosis occurs in fibroblasts and can be induced by various profibrotic stimuli such growth factors or hypoxia, to the best of our knowledge, there are no reports describing a functional importance of this process in pulmonary fibrosis. Thus, we hypothesized that macropinocytosis may contribute to the profibrotic phenotype of activated fibroblasts and to the progression of pulmonary fibrosis. We found that macropinocytosis is upregulated in IPF-derived lung fibroblasts and that inhibition of this process exerts a prominent antifibrotic effect in fibroblasts, Bleo-injured mice and IPF-derived PCLS (**Figures 1, 4, and 5**). Furthermore, we found that macropinocytosis inhibition ameliorates mTORC1 signaling pathway activation by regulating AA uptake in activated lung fibroblasts (**Figure 2**).

A large body of evidence strongly suggests that changes in the cytosolic concentration of AAs are directly sensed by mTORC1 which leads to the activation of various signaling pathways that regulate vital intracellular processes including protein and lipid synthesis, mitochondrial biogenesis and glycolysis (38, 39). mTORC1 has been also shown to contribute to the progression of fibroblast activation and pulmonary fibrosis (26, 27, 40). However, these pre-clinical and experimental findings were challenged when rapamycin, a non-specific mTOR inhibitor, demonstrated limited success as a treatment for IPF compared to placebo in a short-term clinical study (41). Therefore, a precise understanding of mTORC1-mediated profibrotic signaling pathways will be required to develop either more specific mTORC1 inhibitors or those which target profibrotic signaling proteins downstream of mTORC1 activation (42, 43). Recent advances in this area have resulted in the development of a third generation of mTORC1 inhibitors, termed bi-steric mTORC1 inhibitors, which are more specific and effectively regulate downstream signaling proteins, such as S6 kinase (S6K) and 4E-binding protein 1 (4EBP1), with minimal or no effect on mTORC2 (43, 44). These inhibitors have already demonstrated therapeutic effects in experimental models of cancer and lymphangioleiomyomatosis (LAM) (43–45). Since mTORC1 plays a key role in profibrotic responses of fibroblasts, the potential anti-fibrotic activity of bi-steric mTORC1 inhibitors should be also tested in organ fibrosis models in the future. However, the mechanism by which mTORC1 signaling contributes to profibrotic responses in fibroblasts is not fully elucidated.

MEOX1 belongs to a subfamily of antennapedia-like homeobox-containing transcription factors, that recognize a palindromic DNA binding site sequence, TAATTA (46, 47). Early reports have demonstrated that MEOX1 plays critical roles in skeletal and mesenchymal tissue formation during embryogenesis, vascular cell activation and senescence (47–49). Recent reports strongly suggest a pivotal role of MEOX1 in cardiac fibroblast activation and fibrosis (29, 50). More recently, a study conducted by Jin *et al.* has demonstrated that MEOX1 plays a key role in apoptosis resistance in myofibroblasts, a known profibrotic feature of IPF-derived lung fibroblasts. Consistent with previous findings, we found that MEOX1 silencing ameliorated fibroblast activation (**Figure 3 E**). Furthermore, we found that MEOX1 expression is downregulated by RAPTOR silencing or EIPA treatment, which suggests that the induction of MEOX1 expression in fibroblasts is mediated by macropinocytosis followed by mTORC1 activation (**Figure 3 C and D**).

Macropinocytosis is dysregulated in various health disorders, including cancer, and contributes to their pathobiology. Thus, the inhibition of this process may serve as a potential therapeutic strategy. To this end, an elegant study conducted by Lin *et al.,* had screened 640 FDA-approved drugs and identified imipramine, a tricyclic antidepressant, as a leading candidate to serve as a specific inhibitor of macropinocytosis by an unknown mechanism, without affecting phagocytosis, caveolin-or clathrin-mediated endocytosis (21). More recently, another drug screen identified another tricyclic antidepressant, nortriptyline, as a potent macropinocytosis inhibitor with prominent effects in blocking fatty acid uptake and repressing tumor growth. Since repurposing FDA-approved drugs can represent a more efficient and attractive strategy with respect to time and cost compared to *de novo* drug design, we tested the feasibility of imipramine to regulate macropinocytosis and pulmonary fibrosis. As expected, we found that imipramine potently downregulated macropinocytosis, similar to the effects of a known macropinocytosis inhibitor EIPA, without affecting clathrin-dependent endocytosis or caveolin-1 expression (**Supplemental Figure 1**). We also observed that imipramine inhibits lung fibroblast activation and ameliorates fibrosis in mouse and PCLS models of pulmonary fibrosis (**Figure 6**).

Taken together, our results strongly suggest that macropinocytosis inhibition is a feasible therapeutic strategy to ameliorate profibrotic responses in fibroblasts and pulmonary fibrosis in experimental models. We further suggest that repurposing imipramine, an FDA-approved anti-depressant, as a macropinocytosis inhibitor can potentially be considered for future clinical development as an anti-fibrotic therapy.

## METHODS

Further information can be found in Supplemental Methods

### Sex as a biological variable

For human tissue-based studies, both male and female subjects were used. For mouse studies, we examined male and female animals, and similar findings are reported for both sexes.

### Statistical analysis

Data are expressed as mean ± SEM. Comparisons of mortality were made by analyzing Kaplan-Meier survival curves and log-rank tests to assess for differences in survival. For comparisons between two groups, we used Student’s unpaired t test. Statistical significance was defined as *P*<0.05. One-way analysis of variance, followed by Newman-Keuls or Tukey’s post-test analysis was used for analysis of more than two groups. The numbers of samples per group (n), or the numbers of experiments, are specified in the figure legends.

### Study approval

This study was approved by the institutional review board at Baylor College of Medicine to use the deidentified patient specimens (Protocol number: H-50814). All animal experimental protocols were approved by the Baylor College of Medicine Institutional Animal Care and Use Committee (Protocol number: AN-8219).

### Data availability

Data are available in the Supporting Data Values file or associated supplemental files. Bulk and single cell RNA sequencing were deposited in the Gene Expression Omnibus (GEO) database GSE298860, GSE300087 respectively.

## ACKNOWLEDGEMENTS

This work was supported by grants from the National Institute of Arthritis, Musculoskeletal and Skin Diseases (NIAMS) (AR074558), and the National Heart, Lung and Blood Institute (NHLBI) (HL176934), and Veterans Affairs Research Career Scientist award VA IK6 BX005647.

## REFERENCES

1. King TE, Jr., Pardo A, and Selman M. Idiopathic pulmonary fibrosis. Lancet. 2011;378(9807):1949-61.

2. Raghu G, Chen SY, Yeh WS, Maroni B, Li Q, Lee YC, et al. Idiopathic pulmonary fibrosis in US Medicare beneficiaries aged 65 years and older: incidence, prevalence, and survival, 2001-11. Lancet Respir Med. 2014;2(7):566–72.

3. Richeldi L, Costabel U, Selman M, Kim DS, Hansell DM, Nicholson AG, et al. Efficacy of a tyrosine kinase inhibitor in idiopathic pulmonary fibrosis. N Engl J Med. 2011;365(12):1079–87.

4. Richeldi L, du Bois RM, Raghu G, Azuma A, Brown KK, Costabel U, et al. Efficacy and safety of nintedanib in idiopathic pulmonary fibrosis. N Engl J Med. 2014;370(22):2071–82.

5. Pardo A, and Selman M. Lung Fibroblasts, Aging, and Idiopathic Pulmonary Fibrosis. Ann Am Thorac Soc. 2016;13 Suppl 5:S417–S21.

6. White ES, Lazar MH, and Thannickal VJ. Pathogenetic mechanisms in usual interstitial pneumonia/idiopathic pulmonary fibrosis. J Pathol. 2003;201(3):343–54.

7. Thannickal VJ, Toews GB, White ES, Lynch JP, 3rd, and Martinez FJ. Mechanisms of pulmonary fibrosis. Annu Rev Med. 2004;55:395–417.

8. Stow JL, Hung Y, and Wall AA. Macropinocytosis: Insights from immunology and cancer. Curr Opin Cell Biol. 2020;65:131–40.

9. Commisso C, Davidson SM, Soydaner-Azeloglu RG, Parker SJ, Kamphorst JJ, Hackett S, et al. Macropinocytosis of protein is an amino acid supply route in Ras-transformed cells. Nature. 2013;497(7451):633-7.

10. Charpentier JC, Chen D, Lapinski PE, Turner J, Grigorova I, Swanson JA, et al. Macropinocytosis drives T cell growth by sustaining the activation of mTORC1. Nat Commun. 2020;11(1):180.

11. Recouvreux MV, and Commisso C. Macropinocytosis: A Metabolic Adaptation to Nutrient Stress in Cancer. Front Endocrinol (Lausanne). 2017;8:261.

12. Alvarez D, Cardenes N, Sellares J, Bueno M, Corey C, Hanumanthu VS, et al. IPF lung fibroblasts have a senescent phenotype. Am J Physiol Lung Cell Mol Physiol. 2017;313(6):L1164–L73.

13. Geng Y, Liu X, Liang J, Habiel DM, Kulur V, Coelho AL, et al. PD-L1 on invasive fibroblasts drives fibrosis in a humanized model of idiopathic pulmonary fibrosis. JCI Insight. 2019;4(6).

14. Tsoyi K, Liang X, De Rossi G, Ryter SW, Xiong K, Chu SG, et al. CD148 Deficiency in Fibroblasts Promotes the Development of Pulmonary Fibrosis. Am J Respir Crit Care Med. 2021.

15. Commisso C, Flinn RJ, and Bar-Sagi D. Determining the macropinocytic index of cells through a quantitative image-based assay. Nat Protoc. 2014;9(1):182–92.

16. Lee SW, Zhang Y, Jung M, Cruz N, Alas B, and Commisso C. EGFR-Pak Signaling Selectively Regulates Glutamine Deprivation-Induced Macropinocytosis. Dev Cell. 2019;50(3):381–92 e5.

17. Zhang MS, Cui JD, Lee D, Yuen VW, Chiu DK, Goh CC, et al. Hypoxia-induced macropinocytosis represents a metabolic route for liver cancer. Nat Commun. 2022;13(1):954.

18. Koivusalo M, Welch C, Hayashi H, Scott CC, Kim M, Alexander T, et al. Amiloride inhibits macropinocytosis by lowering submembranous pH and preventing Rac1 and Cdc42 signaling. J Cell Biol. 2010;188(4):547–63.

19. Lin XP, Mintern JD, and Gleeson PA. Macropinocytosis in Different Cell Types: Similarities and Differences. Membranes (Basel). 2020;10(8).

20. Ivanov AI. Pharmacological inhibition of endocytic pathways: is it specific enough to be useful? Methods Mol Biol. 2008;440:15–33.

21. Lin HP, Singla B, Ghoshal P, Faulkner JL, Cherian-Shaw M, O’Connor PM, et al. Identification of novel macropinocytosis inhibitors using a rational screen of Food and Drug Administration-approved drugs. Br J Pharmacol. 2018;175(18):3640–55.

22. Nagano T, Iwasaki T, Onishi K, Awai Y, Terachi A, Kuwaba S, et al. LY6D-induced macropinocytosis as a survival mechanism of senescent cells. J Biol Chem. 2020.

23. Seguin L, Odouard S, Corlazzoli F, Haddad SA, Moindrot L, Calvo Tardon M, et al. Macropinocytosis requires Gal-3 in a subset of patient-derived glioblastoma stem cells. Commun Biol. 2021;4(1):718.

24. Yoshida S, Pacitto R, Yao Y, Inoki K, and Swanson JA. Growth factor signaling to mTORC1 by amino acid-laden macropinosomes. J Cell Biol. 2015;211(1):159–72.

25. Zhang Y, Recouvreux MV, Jung M, Galenkamp KMO, Li Y, Zagnitko O, et al. Macropinocytosis in Cancer-Associated Fibroblasts Is Dependent on CaMKK2/ARHGEF2 Signaling and Functions to Support Tumor and Stromal Cell Fitness. Cancer Discov. 2021;11(7):1808–25.

26. Hu X, Zhang H, Li X, Li Y, and Chen Z. Activation of mTORC1 in fibroblasts accelerates wound healing and induces fibrosis in mice. Wound Repair Regen. 2020;28(1):6–15.

27. Selvarajah B, Azuelos I, Plate M, Guillotin D, Forty EJ, Contento G, et al. mTORC1 amplifies the ATF4-dependent de novo serine-glycine pathway to supply glycine during TGF-beta1-induced collagen biosynthesis. Sci Signal. 2019;12(582).

28. Woodcock HV, Eley JD, Guillotin D, Plate M, Nanthakumar CB, Martufi M, et al. The mTORC1/4E-BP1 axis represents a critical signaling node during fibrogenesis. Nat Commun. 2019;10(1):6.

29. Schumacher D, Peisker F, and Kramann R. MEOX1: a novel druggable target that orchestrates the activation of fibroblasts in cardiac fibrosis. Signal Transduct Target Ther. 2021;6(1):440.

30. Lin HP, Singla B, Ahn W, Ghoshal P, Blahove M, Cherian-Shaw M, et al. Receptor-independent fluid-phase macropinocytosis promotes arterial foam cell formation and atherosclerosis. Sci Transl Med. 2022;14(663):eadd2376.

31. Tsukui T, Wolters PJ, and Sheppard D. Alveolar fibroblast lineage orchestrates lung inflammation and fibrosis. Nature. 2024;631(8021):627-34.

32. Celada SI, Lim CX, Carisey AF, Ochsner SA, Arce Deza CF, Rexie P, et al. SHP2 promotes sarcoidosis severity by inhibiting SKP2-targeted ubiquitination of TBET in CD8(+) T cells. Sci Transl Med. 2023;15(713):eade2581.

33. Ochsner SA, Pillich RT, and McKenna NJ. Consensus transcriptional regulatory networks of coronavirus-infected human cells. Sci Data. 2020;7(1):314.

34. Bar-Sagi D, and Feramisco JR. Induction of membrane ruffling and fluid-phase pinocytosis in quiescent fibroblasts by ras proteins. Science. 1986;233(4768):1061-8.

35. Veithen A, Cupers P, Baudhuin P, and Courtoy PJ. v-Src induces constitutive macropinocytosis in rat fibroblasts. J Cell Sci. 1996;109 ( Pt 8):2005–12.

36. Mellstrom K, Heldin CH, and Westermark B. Induction of circular membrane ruffling on human fibroblasts by platelet-derived growth factor. Exp Cell Res. 1988;177(2):347–59.

37. Zhang YF, Li Q, Huang PQ, Su T, Jiang SH, Hu LP, et al. A low amino acid environment promotes cell macropinocytosis through the YY1-FGD6 axis in Ras-mutant pancreatic ductal adenocarcinoma. Oncogene. 2022;41(8):1203–15.

38. Saxton RA, and Sabatini DM. mTOR Signaling in Growth, Metabolism, and Disease. Cell. 2017;168(6):960–76.

39. Zoncu R, Bar-Peled L, Efeyan A, Wang S, Sancak Y, and Sabatini DM. mTORC1 senses lysosomal amino acids through an inside-out mechanism that requires the vacuolar H(+)-ATPase. Science. 2011;334(6056):678-83.

40. O’Leary EM, Tian Y, Nigdelioglu R, Witt LJ, Cetin-Atalay R, Meliton AY, et al. TGF-beta Promotes Metabolic Reprogramming in Lung Fibroblasts via mTORC1-dependent ATF4 Activation. Am J Respir Cell Mol Biol. 2020;63(5):601–12.

41. Gomez-Manjarres DC, Axell-House DB, Patel DC, Odackal J, Yu V, Burdick MD, et al. Sirolimus suppresses circulating fibrocytes in idiopathic pulmonary fibrosis in a randomized controlled crossover trial. JCI Insight. 2023;8(8).

42. Yan XL, Pan YH, Fan RZ, Song QQ, Li S, Huang JL, et al. Discovery of the First Raptor (Regulatory-Associated Protein of mTOR) Inhibitor as a New Type of Antiadipogenic Agent. J Med Chem. 2023;66(8):5839–58.

43. Burnett GL, Yang YC, Aggen JB, Pitzen J, Gliedt MK, Semko CM, et al. Discovery of RMC-5552, a Selective Bi-Steric Inhibitor of mTORC1, for the Treatment of mTORC1-Activated Tumors. J Med Chem. 2023;66(1):149–69.

44. Du H, Yang YC, Liu HJ, Yuan M, Asara JM, Wong KK, et al. Bi-steric mTORC1 inhibitors induce apoptotic cell death in tumor models with hyperactivated mTORC1. J Clin Invest. 2023;133(21).

45. Evans JF, Ledwell OA, Tang Y, Rue R, Mukhitov AR, Diesler R, et al. The Bi-Steric Inhibitor RMC-5552 Reduces mTORC1 Signaling and Growth in Lymphangioleiomyomatosis. Am J Respir Cell Mol Biol. 2024.

46. Candia AF, Hu J, Crosby J, Lalley PA, Noden D, Nadeau JH, et al. Mox-1 and Mox-2 define a novel homeobox gene subfamily and are differentially expressed during early mesodermal patterning in mouse embryos. Development. 1992;116(4):1123–36.

47. Skuntz S, Mankoo B, Nguyen MT, Hustert E, Nakayama A, Tournier-Lasserve E, et al. Lack of the mesodermal homeodomain protein MEOX1 disrupts sclerotome polarity and leads to a remodeling of the cranio-cervical joints of the axial skeleton. Dev Biol. 2009;332(2):383–95.

48. Douville JM, Cheung DY, Herbert KL, Moffatt T, and Wigle JT. Mechanisms of MEOX1 and MEOX2 regulation of the cyclin dependent kinase inhibitors p21 and p16 in vascular endothelial cells. PLoS One. 2011;6(12):e29099.

49. Rodrigo I, Bovolenta P, Mankoo BS, and Imai K. Meox homeodomain proteins are required for Bapx1 expression in the sclerotome and activate its transcription by direct binding to its promoter. Mol Cell Biol. 2004;24(7):2757–66.

50. Alexanian M, Przytycki PF, Micheletti R, Padmanabhan A, Ye L, Travers JG, et al. A transcriptional switch governs fibroblast activation in heart disease. Nature. 2021;595(7867):438-43.

