## Supplementary material for "Macropinocytosis inhibition attenuates pro-fibrotic responses in lung fibroblasts and pulmonary fibrosis": Suplemental Information

**Methods:**

**Reagents**

Antibodies against α-SMA (Cat #: ab5694), collagen 1 (Cat #: ab316222), and fibronectin (Cat #: ab2413) were from Abcam. Anti-GAPDH, and anti-MEOX1 antibodies were obtained from Santa Cruz Biotechnology (Cat#: sc-365062, and sc-398845 respectively). Antibodies against phospho-S6 ribosomal protein (Ser235/236, Cat #: 2211), S6 ribosomal protein (Cat #: 2217), phospho-p70 S6 kinase (Thr389, Cat #: 9206), p70 S6 kinase (Cat #: 2708) were from Cell Signaling Technologies. Recombinant TGF-β1 was obtained from Sino Biological (Cat. # 10804-HNAC). Rapalink-1 was obtained from Cell Signaling Technology (Cat#: 88626). Fluorescein isothiocyanate (FITC)-labeled dextran (70 kDa, Cat #: 46945) was obtained Millipore Sigma. Alexa Fluor-488 (AF488)-labeled transferrin from human Serum was from ThermoFisher Scientific. 5-(N-ethyl-N-isopropyl)-Amiloride (EIPA) (Cat #: 14406), and Imipramine (Cat #: 15890) were obtained from Cayman Chemical. All other reagents, including 4-hydroxy-tamoxifen (4-OHT, Cat #: H6278), were from Millipore Sigma.

**Primary Lung Fibroblasts**

Deidentified human lung fibroblasts (HLFs) were derived from patients who were subjected to lung transplantation with IPF. Control (Con) HLFs were derived from nonfibrotic lung samples lacking any evidence of disease which were unsuitable for transplantation. Mouse lung fibroblasts from *Slc9a1^fl/fl^* Col1a2-Cre-ER(T)^+/0^, *Slc9a1^fl/fl^* mice were obtained as previously described (1). Fibroblasts were cultured in complete media (DMEM; Corning) containing 10% FBS (Corning), 100 IU of penicillin and 100 μg/ml streptomycin (Corning), 292 μg/ml L-glutamine (Corning), and 100 μg/ml Primocin (InVivoGen) in humidified incubators at 37°C and 10% CO_2_. To induce Cre recombinase expression in *Slc9a1^fl/fl^* / Col1a2-Cre-ER(T)^+/0^ or *Slc9a1^fl/fl^* lung fibroblasts, cells were treated with 4-hydroxytamoxifen (4-OHT, 1 μM) for 24 hours.

**Macropinocytosis assay**

Macropinocytosis levels were measured by the uptake of high molecular weight (70 kDa) FITC-labeled dextran by fibroblasts as described with modifications (2). Briefly, cells were cultured on 6 well plate (flow cytometry) or on the slide chamber in complete media. When cells reached 70-80% confluence, they were conditioned in low serum conditions (1% of FBS) for 12-16 hours. Then, cells were treated with 70 kDa FITC-labeled dextran (0.5 mg/ml) for 1 hour. After incubation cells were washed three times with PBS and either fixed with 4% paraformaldehyde, additionally stained with DAPI for fluorescent microscopy imaging or harvested, stained with DAPI and subjected to flow cytometry. Images were analyzed by ImageJ (Image Processing and Analysis in Java Edition: 1.29 URL: http://rsb.info.nih.gov/ij/ NIH, Maryland) and flow cytometry data by FlowJo.

**Mice**

WT C57BL/6 mice were obtained from JAX laboratory and used at 10-12 weeks of age. Conditional Slc9a1-knockout mice of both sexes were bred as follows on a C57BL/6 background. Transgenic Col1a2^Cre-ER(T)+/0^ mice were obtained from JAX Laboratory (Strain #: 029567) and *Slc9a1^fl/fl^* mice were donated by Dr. Dandan Sun (University of Pittsburgh) (3). To generate fibroblast-specific Slc9a1-deficient mice, *Slc9a1^fl/fl^* mice were bred with Col1a2^Cre-ER(T)+/0^ (heterozygous allele) transgenic mice to generate mice heterozygous for both alleles. Progeny from the second cross between *Slc9a1^fl/fl^* mice and heterozygous *Slc9a1^fl/WT^* Col1a2^Cre-ER(T)+/0^ mice (from the first cross) were used for further experiments. All mice were genotyped by PCR techniques as described previously (3). For treatment of mice, a stock solution of tamoxifen was diluted in corn oil to 20 mg/ml. To selectively delete Slc9a1 in activated fibroblasts, adult *Slc9a1^fl/fl^* Col1a2^Cre-ER(T)+/0^ mice (10-12 weeks old) and control *Slc9a1^fl/fl^* mice were administered a tamoxifen suspension (0.1 ml of diluted stock) *via* intraperitoneal (*i.p.*) injection (75 mg/kg), for 5 days before administration of bleomycin sulfate (Bleo) (0.5 mg/kg) and every 5 days thereafter until sacrifice.

**Bleomycin (Bleo) model of pulmonary fibrosis**

Lung fibrosis was elicited in mice by intratracheal (*i.t.*) injection of a single dose of 0.5 mg/kg body weight of Bleo as described previously (4); control mice received a volume of sterile saline equal to the volume of Bleo. Mice were sacrificed 21 days after saline or Bleo instillation. Bronchoalveolar lavage fluid (BALF) was collected by lavaging lungs with 1 ml of PBS. BALF was used for protein concentration using bicinchoninic acid (BCA) protein assay, and monocyte chemoattractant protein-1 (MCP1) by ELISA. The right middle lobe was subjected to hydroxyproline assay, the left lung was fixed and subjected to immunohistochemistry, the rest of the right lung was assessed for expression of fibrotic marker mRNAs (*Acta2*, *Col1a1*, and *Meox1*) and histology.

**Hydroxyproline assay**

To quantify collagen deposition, the right lung (middle lobe) from each mouse was hydrolyzed in 6N HCl for 24 h at 110°C, and hydroxyproline levels were quantified as previously described (5). Each sample was tested in triplicate. Data are expressed as micrograms of hydroxyproline per lung.

**Histology**

Lung sections were fixed by inflation with buffered 10% formalin solution and embedded in paraffin. Thin (4 μm) sections were deparaffinized and rehydrated. Tissue slides were then stained by hematoxylin-eosin (H&E) and Masson's Trichrome staining to evaluate histopathologic changes and collagen deposition, respectively (5).

**Precision Cut Lung Slices (PCLS)**

PCLS from IPF lungs were prepared as previously described. Briefly, using a syringe pump, lungs were infiltrated with warm, 2% (37°C) low-melting agarose–HBSS solution (MilliporeSigma, no. A9414; kept at 37°C). After complete solidification of agarose in the inflated lobes on ice, tissue blocks of approximately 10 mm in diameter were prepared. Lung slices (300 μm thick) were cut perpendicularly to the visible airway with a vibratome (Precisionary Instruments, no. VF-300, Greenville, NC) at room temperature in HBSS. Then, slices were cultured in 24-well plates supplemented with DMEM/F12 media containing 1% FBS and antibiotics. Lung slices were incubated with or without EIPA (12.5 μM) or Imipramine (Imi, 10 μM) for another 72 h. Culture media was changed at 24 and 48 hours and treated with drugs or DMSO with total of 2 treatments. After treatment, slices were subjected to total RNA isolation to measure profibrotic genes expression (*ACTA2, COL1A1*) or immunofluorescent staining.

**Single Cell RNA sequencing**

PCLS from IPF lungs were treated Single cell lung data from was mapped using 10x Genomics Cell Ranger v7.0.1 onto the human genome reference provided by 10x Genomics. Doublet detection was performed using scrubblet (6). Data was processed using the Python Scanpy library (7) we performed guided cell annotation using CellTypist (8) and references provided by the Human Lung Cell Atlas (HLCA) (9) augmented with aberrant basaloid reference (10). UMAP plots were generated using Scanpy. Objects were converted to the Seurat format using the zellkonverter package. Cell type marker plots were generated using Seurat (11). Differential gene expression was generated using MAST (12) with significance achieved at FDR-adjusted p-value<0.1 and fold change exceeding 1.25x. Relative differences in abundance of cell types within specific compartments were derived using ChiSquare as implemented in the R statistical system.

**Bulk RNA sequencing**

MRC-5 cells were treated with EIPA (12.5 μM) or Vehicle (DMSO) with or without TGF-β1 (10 ng/ml) for 24 hours. Then total RNA isolation and elimination of DNA was done using RNeasy kit (QIAGEN). A cDNA library was prepared with 250ng of RNA using the Illumina TruSeq Stranded mRNA kit according to the manufacturer’s protocol. Libraries were quantitated and pooled equimolarly. Sequencing was performed on an Illumina NextSeq 500 targeting around 40 million read pairs per sample. Sequencing was run on NextSeq 500 Sequencing System. FastQ file generation was executed using Illumina’s cloud-based informatics platform, BaseSpace Sequencing Hub at BCM Genomic and RNA Profiling Core.

**Western Blot**

Polyacrylamide gel electrophoresis and immunoblotting were performed according to standard methods as previously described (13). The electrophoresed proteins were transferred to polyvinylidene difluoride (PVDF) membranes by semidry electrophoretic transfer at 15 V for 60–75 min. The membranes were blocked overnight at 4°C in 5% bovine serum albumin (BSA). The cells were incubated with primary antibodies diluted 1:500 in Tris-buffered saline/Tween 20 (TBS-T) containing 5% BSA for 2 h and then incubated with the secondary antibody at room temperature for 1 h. Suitable horseradish peroxidase (HRP)-conjugated secondary antibodies were used (1:5000 dilution in TBST containing 1% BSA). The signals were detected by ECL (Thermo Fisher Scientific). Quantification of protein bands was performed with the computer software ImageJ and was expressed as a ratio of band intensity with respect to the loading control.

**Quantitative Real-Time PCR (qPCR)**

Total RNA was isolated with Trizol reagent (Invitrogen, Carlsbad, CA) and cDNA was synthesized from RNA (1 μg) using a SuperScript First-strand synthesis system for reverse transcription (RT) PCR (Invitrogen, Carlsbad, CA). Primer sequences are shown in Table S1. RT-PCR, with SYBR Green Master Mix (Bio-Rad Laboratories, Hercules, CA, USA), was performed using the StepOnePlus Real-Time PCR System (Applied Biosystems). The relative quantity of target mRNA was calculated by use of the CT method, or 2–∆∆CT, as described (14), and normalized by use of GAPDH as an endogenous control (Sequence Detection System software, version 1.7; Applied Biosystems).

**Amino acid (AA)-free and -supplemented conditions and amino acid assay**

Cells were plated on 24 well plates at a density of 1 x 10^4^ per well. Then, cells were incubated in AA free and AA-supplemented DMEM media. AA-free media was prepared from commercially available AA-free DMEM powder (MyBiosource, Cat #: MBS6120661) and supplemented with antibiotics or supplemented with MEM amino acid solution (ThermoFisher Scientific, Cat #: 11130051) and glutamine (referred as AA-supplemented media) in the presence or absence of EIPA (12.5 μM) ± TGF-β1 (10 ng/ml) for 4 hours. Then, cells were washed with PBS twice, lysed and subjected to L-amino acid assay using a commercially available kit (Abcam, Cat #: ab65347).

**Gel contraction assay**

The assay was performed as previously described (5). Gel sizes were measured by the ruler at 0 and 24 h.

**Mitochondrial stress assay**

HLFs were treated with EIPA (12.5 μM) ± TGF-β1 (10 ng/ml) for 24 hours. Then, cells were replated onto Seahorse XFe24 plates at 3 × 10^4^ per well, cultured in Seahorse XF DMEM medium, pH 7.4 (Agilent, Cat. # 103575-100). Media was supplemented with 10 mM glucose, 2 mM glutamine, and 1 mM pyruvate. Injections of oligomycin A (1 μM), carbonyl cyanide-4-trifluoromethoxyphenylhydrazone (FCCP, 2 μM), and antimycin A and rotenone (1 μM) supplied by Agilent (Seahorse XFp Cell Mito Stress Test Kit, Cat #: 103010-100). Oxygen consumption rate (OCR) was detected using Seahorse XFe24 Analyzer.

**Lentiviral Transfection**

For TSC1, RAPTOR, MEOX1 silencing experiments, the pLKO.1 plasmid, carrying the human shRNA against target genes (consortium numbers TRCN0000010453, TRCN0000010415, TRCN0000016110 respectively) and pLKO.1, carrying a Scr sequence, were purchased from MiliporeSigma. Lentiviral particles were generated by use of a commercially available packaging mix, provided by Millipore Sigma (Cat. #: SHP001) in human embryonic kidney 293 T cells, according to the manufacturer's instructions. Lung fibroblasts were infected with the lentiviral particles, and stably infected cells were selected by use of puromycin (10 μg/ml).

**Cell viability**

Cell viability was determined using the 3‐[4,5-dimethylthiazol‐2‐yl]-2,5-diphenyl tetrazolium bromide (MTT) assay as previously described (15).

**References:**

1.	Tsoyi K, Hall SR, Dalli J, Colas RA, Ghanta S, Ith B, et al. Carbon Monoxide Improves Efficacy of Mesenchymal Stromal Cells During Sepsis by Production of Specialized Proresolving Lipid Mediators. *Crit Care Med.* 2016;44(12):e1236-e45.

2.	Commisso C, Flinn RJ, and Bar-Sagi D. Determining the macropinocytic index of cells through a quantitative image-based assay. *Nat Protoc.* 2014;9(1):182-92.

3.	Begum G, Song S, Wang S, Zhao H, Bhuiyan MIH, Li E, et al. Selective knockout of astrocytic Na(+) /H(+) exchanger isoform 1 reduces astrogliosis, BBB damage, infarction, and improves neurological function after ischemic stroke. *Glia.* 2018;66(1):126-44.

4.	Tsoyi K, Liang X, De Rossi G, Ryter SW, Xiong K, Chu SG, et al. CD148 Deficiency in Fibroblasts Promotes the Development of Pulmonary Fibrosis. *Am J Respir Crit Care Med.* 2021.

5.	Tsoyi K, Chu SG, Patino-Jaramillo NG, Wilder J, Villalba J, Doyle-Eisele M, et al. Syndecan-2 Attenuates Radiation-induced Pulmonary Fibrosis and Inhibits Fibroblast Activation by Regulating PI3K/Akt/ROCK Pathway via CD148. *Am J Respir Cell Mol Biol.* 2018;58(2):208-15.

6.	Wolock SL, Lopez R, and Klein AM. Scrublet: Computational Identification of Cell Doublets in Single-Cell Transcriptomic Data. *Cell Syst.* 2019;8(4):281-91 e9.

7.	Wolf FA, Angerer P, and Theis FJ. SCANPY: large-scale single-cell gene expression data analysis. *Genome Biol.* 2018;19(1):15.

8.	Xu C, Prete M, Webb S, Jardine L, Stewart BJ, Hoo R, et al. Automatic cell-type harmonization and integration across Human Cell Atlas datasets. *Cell.* 2023;186(26):5876-91 e20.

9.	Sikkema L, Ramirez-Suastegui C, Strobl DC, Gillett TE, Zappia L, Madissoon E, et al. An integrated cell atlas of the lung in health and disease. *Nat Med.* 2023;29(6):1563-77.

10.	Adams TS, Schupp JC, Poli S, Ayaub EA, Neumark N, Ahangari F, et al. Single-cell RNA-seq reveals ectopic and aberrant lung-resident cell populations in idiopathic pulmonary fibrosis. *Sci Adv.* 2020;6(28):eaba1983.

11.	Stuart T, Butler A, Hoffman P, Hafemeister C, Papalexi E, Mauck WM, 3rd, et al. Comprehensive Integration of Single-Cell Data. *Cell.* 2019;177(7):1888-902 e21.

12.	Finak G, McDavid A, Yajima M, Deng J, Gersuk V, Shalek AK, et al. MAST: a flexible statistical framework for assessing transcriptional changes and characterizing heterogeneity in single-cell RNA sequencing data. *Genome Biol.* 2015;16:278.

13.	Tsoyi K, Lee TY, Lee YS, Kim HJ, Seo HG, Lee JH, et al. Heme-oxygenase-1 induction and carbon monoxide-releasing molecule inhibit lipopolysaccharide (LPS)-induced high-mobility group box 1 release in vitro and improve survival of mice in LPS- and cecal ligation and puncture-induced sepsis model in vivo. *Mol Pharmacol.* 2009;76(1):173-82.

14.	Schmittgen TD, and Livak KJ. Analyzing real-time PCR data by the comparative C(T) method. *Nat Protoc.* 2008;3(6):1101-8.

15.	Tsoyi K, Kim HJ, Shin JS, Kim DH, Cho HJ, Lee SS, et al. HO-1 and JAK-2/STAT-1 signals are involved in preferential inhibition of iNOS over COX-2 gene expression by newly synthesized tetrahydroisoquinoline alkaloid, CKD712, in cells activated with lipopolysacchride. *Cell Signal.* 2008;20(10):1839-47.

**Figure legends:**

**Supplemental Figure 1. EIPA and Imipramine inhibit macropinocytosis but not other types of endocytosis.**

(A) Human control (Con) and IPF-derived lung fibroblasts were treated with EIPA (12.5 μM) and imipramine (Imi, 10 μM) for 24 hours. After incubation culture media was changed and cells were incubated with FITC-Dextran (70 kDa, 0.5 mg/ml) for 1 hour. Then, cells were washed three times with PBS and subjected to flow cytometry to measure fluorescence intensity of FITC in cells (n=4 each condition). (B) Cells were treated with EIPA and Imi as described in (A). After incubation, cells were exposed to FITC-labeled transferrin (1 mg/ml) for 1 hour. Then cells were washed, harvested and subjected to flow cytometry (n=4 each condition). (C) Cells were treated with EIPA and Imi as described in (A). After treatment cells were lysed and subjected to western blot to determine caveolin-1 (CAV1) (n=4 each condition). *P* < 0.05; significant comparisons by one-way ANOVA: *vs. Control HLF or unstimulated, ^†^vs. IPF HLF.

**Supplemental Figure 2: Effect of EIPA on metabolic reprogramming and cell viability.**

(A) Control HLFs were treated with EIPA (12.5 μM) with or without TGF-β1 (10 ng/ml) for 24 hours. Then cells were replated to Seahorse plates and subjected to the mito-stress assay as described in Methods (n=5 each condition). (B) Control HLFs were treated with EIPA (12.5, and 25 μM) or imipramine (Imi, 10 and 20 μM) for 24 hours. Then, cell viability was measured using MTT assay as described in Methods (n=4 each condition). (C and D) HLFs were lentivirally-transfected with scramble (Scr) or shTSC1 as described in Methods. Then, total RNA was isolated and mRNA levels of *MEOX1* and *TSC1* were determined by RT-qPCR (n=3 or 4 each condition). (E and F) Cells were lentivirally-transfected with scramble (Scr), shRAPTOR (E) or shMEOX1 (F) as described in Methods. Target gene silencing efficiency was measured by RT-qPCR as described in Methods (n=4 each condition). *P* < 0.05; significant comparisons by Student t-test or one-way ANOVA: *vs. unstimulated or Scr, ^†^vs. TGF-β1 alone.

**Supplemental Figure 3: Effect of Slc9a1 deficiency on macropinocytosis in mouse lung fibroblasts.**

Mouse lung fibroblasts (MLFs) were isolated from *Slc9a1^fl/fl^* and *Slc9a1^fl/fl^* Col1a2^Cre-ER(T)+/0^ mice as described in Methods. (A) Slc9a1 mRNA expression was determined with qRT-PCR (n=3 each condition). (B, C and D) Cells were subjected to low AA conditions for 24 hours. Then, cells were incubated with 70kDa FITC-Dextran (0.5 mg/ml) (B), FITC-Transferrin (1 mg/ml) (C), or lysed for western blot to measure CAV1 (n=3 each condition). *P* < 0.05; significant comparisons by Student t-test:* vs. *Slc9a1^fl/fl^* (WT).

**Supplemental Figure 4: H&E staining of the Bleo injured lungs.**

(A and B) Mouse lungs were harvested and stained with H&E on the same area shown for Trichrome staining in Fig. 4 A and C, as described in Methods.

**Supplemental Figure 5: Macropinocytosis inhibitors do not affect lung inflammation in Bleo-injured mice.**

(A) Mouse lungs were stained with H&E on the same area shown for Trichrome staining in Fig. 6 D. B and C. BALF fluid samples were collected as described in Methods and total protein levels and MCP1 were measured by BCA assay or ELISA respectively (n=5 each condition). *P* < 0.05; significant comparisons by Student t-test or one-way ANOVA: *vs. No Bleo (Sham), ^†^vs. Bleo alone.

**Supplemental Figure 6: The effect of EIPA on different mesenchymal cell populations in IPF PCLS.** IPF-derived PCLS were treated with EIPA (12.5 μM) as described in Methods.

**Table 1: List of Primers used for qRT-PCR.**

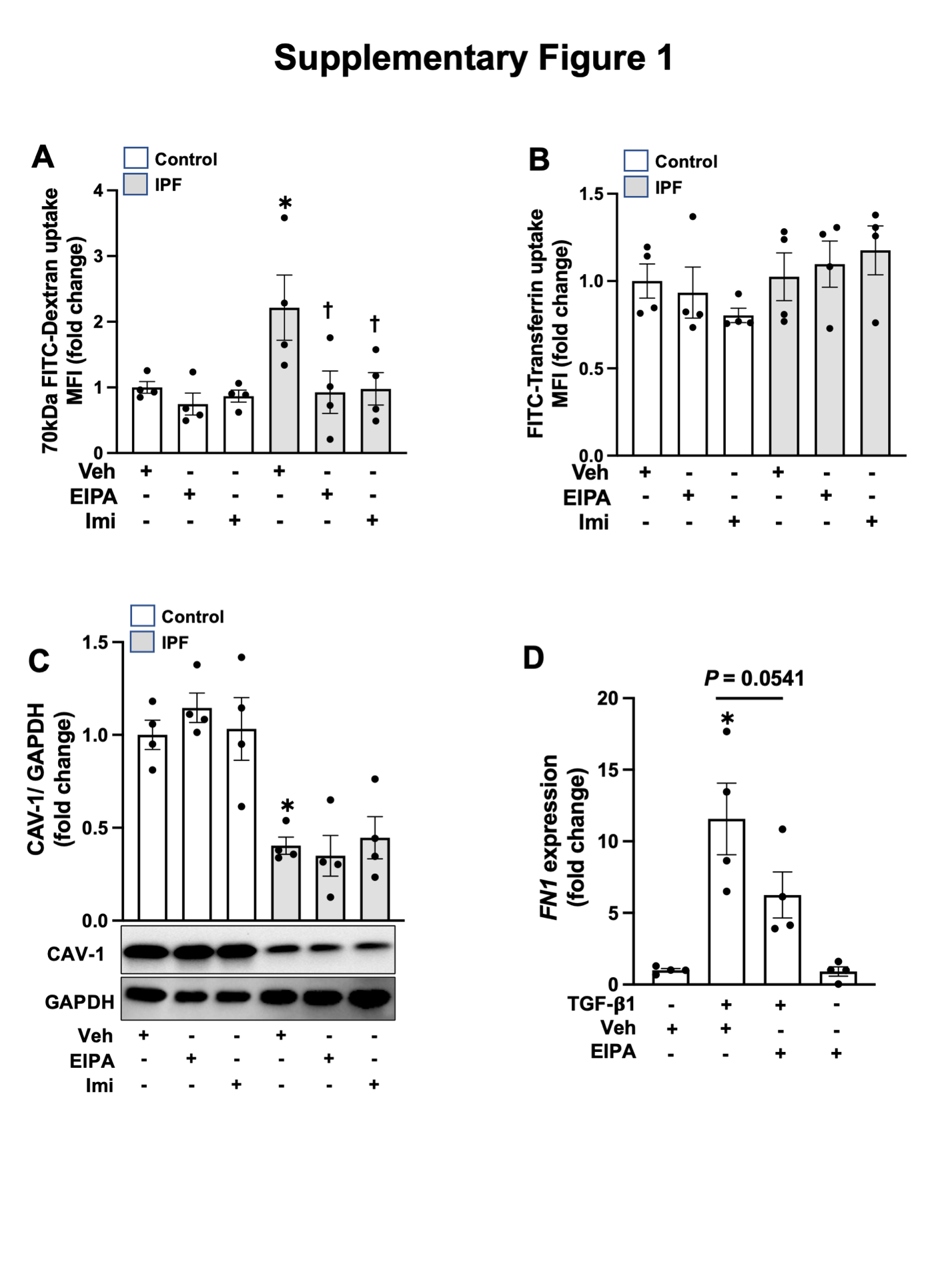

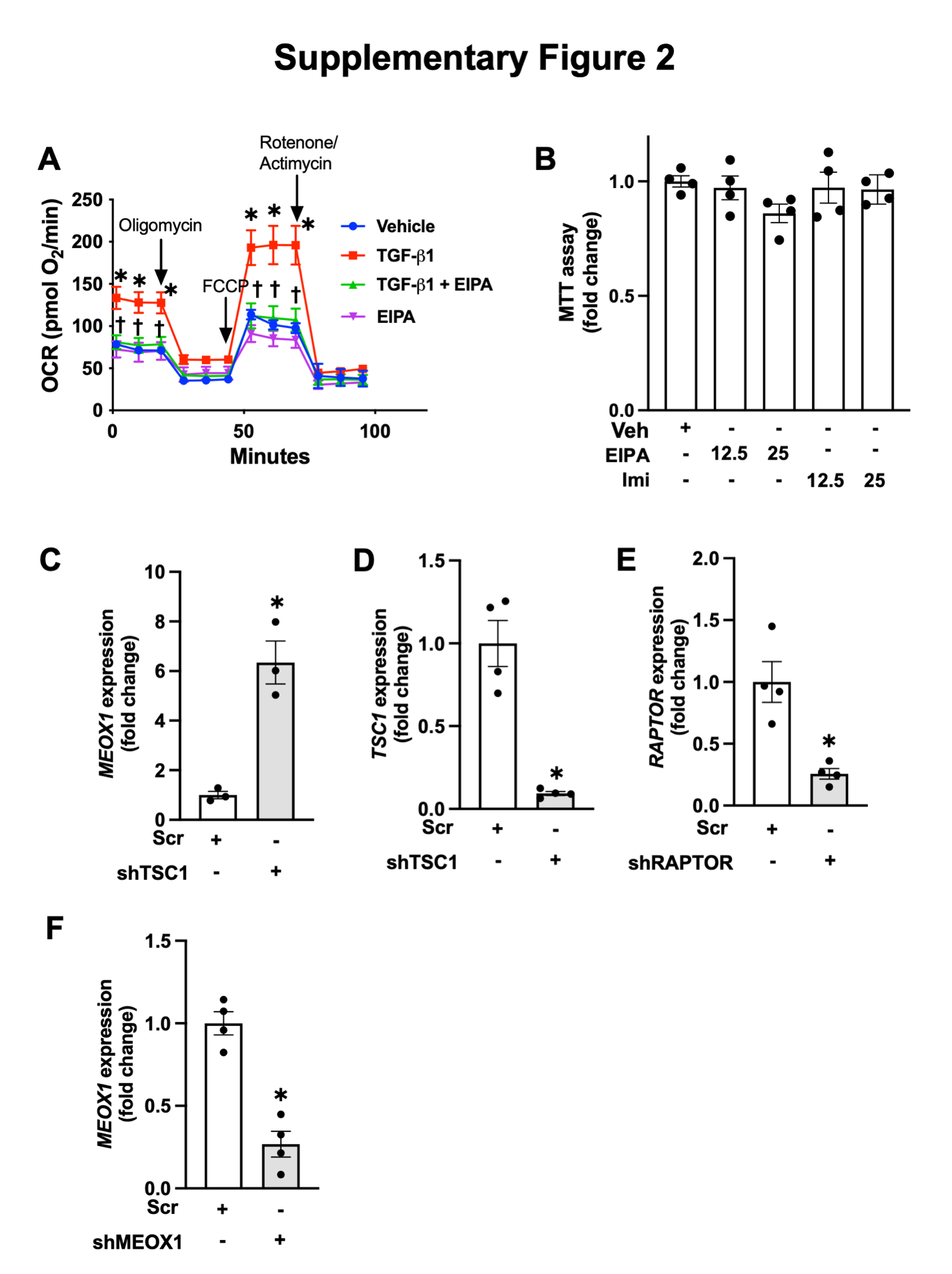

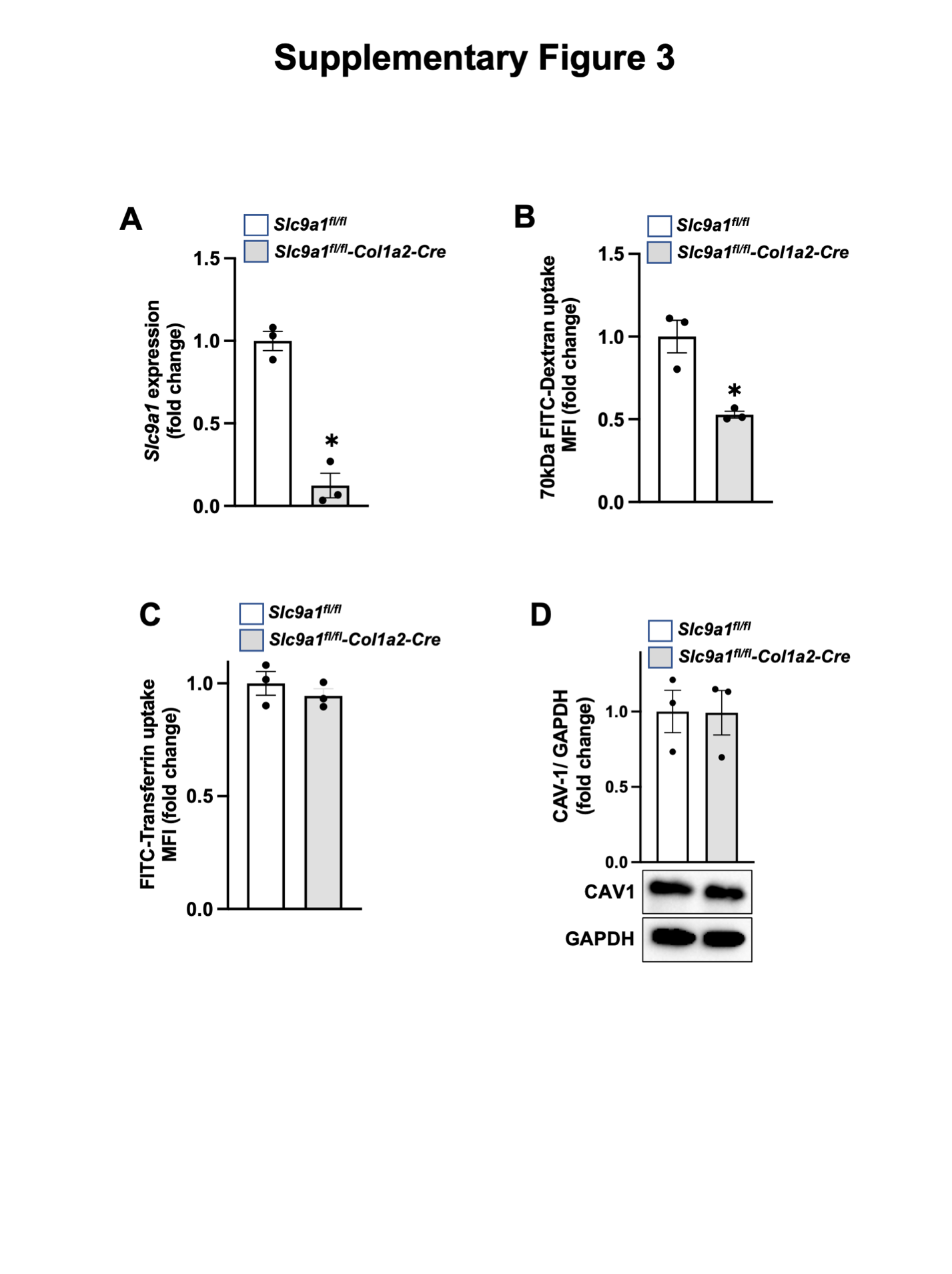

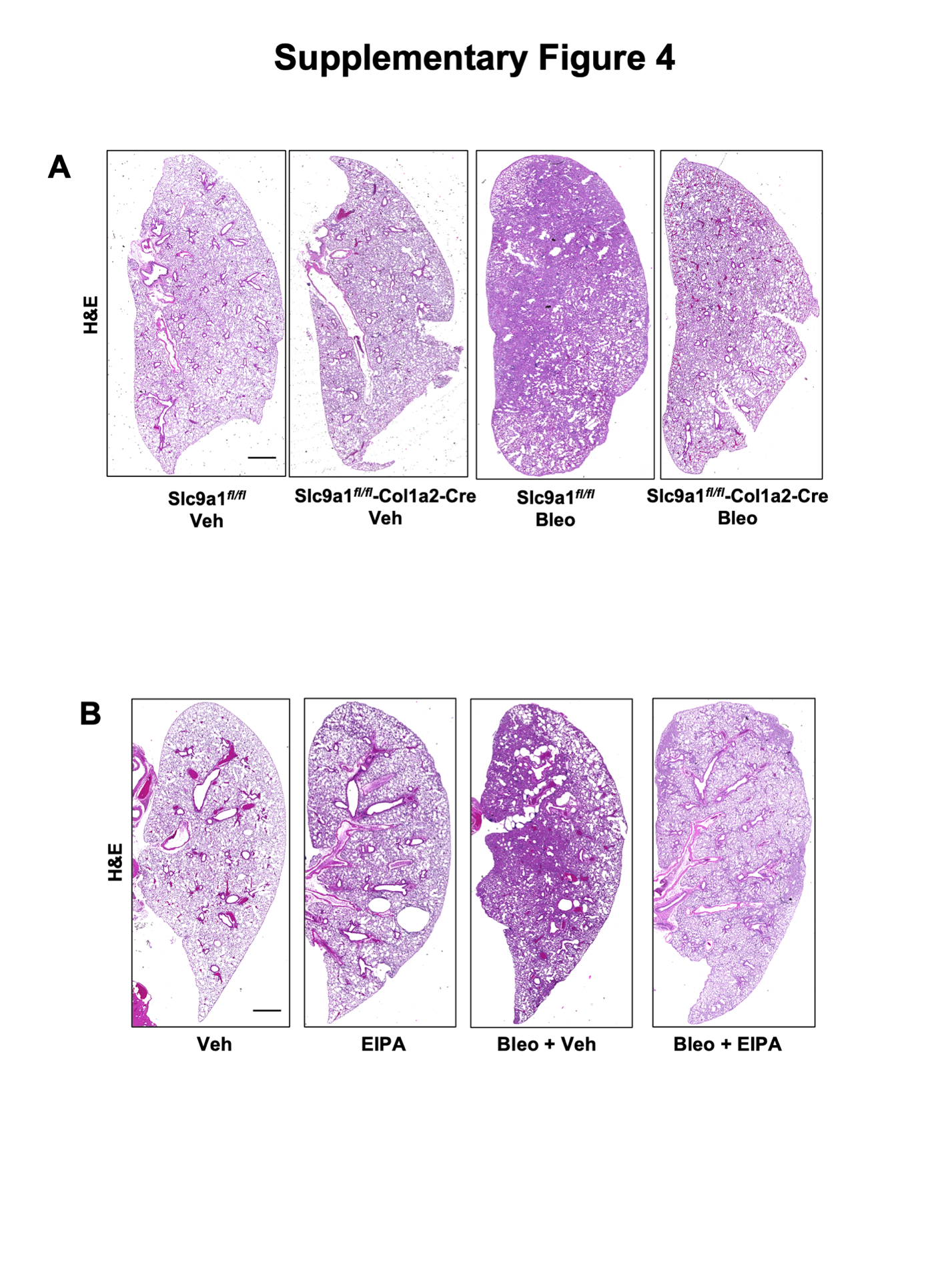

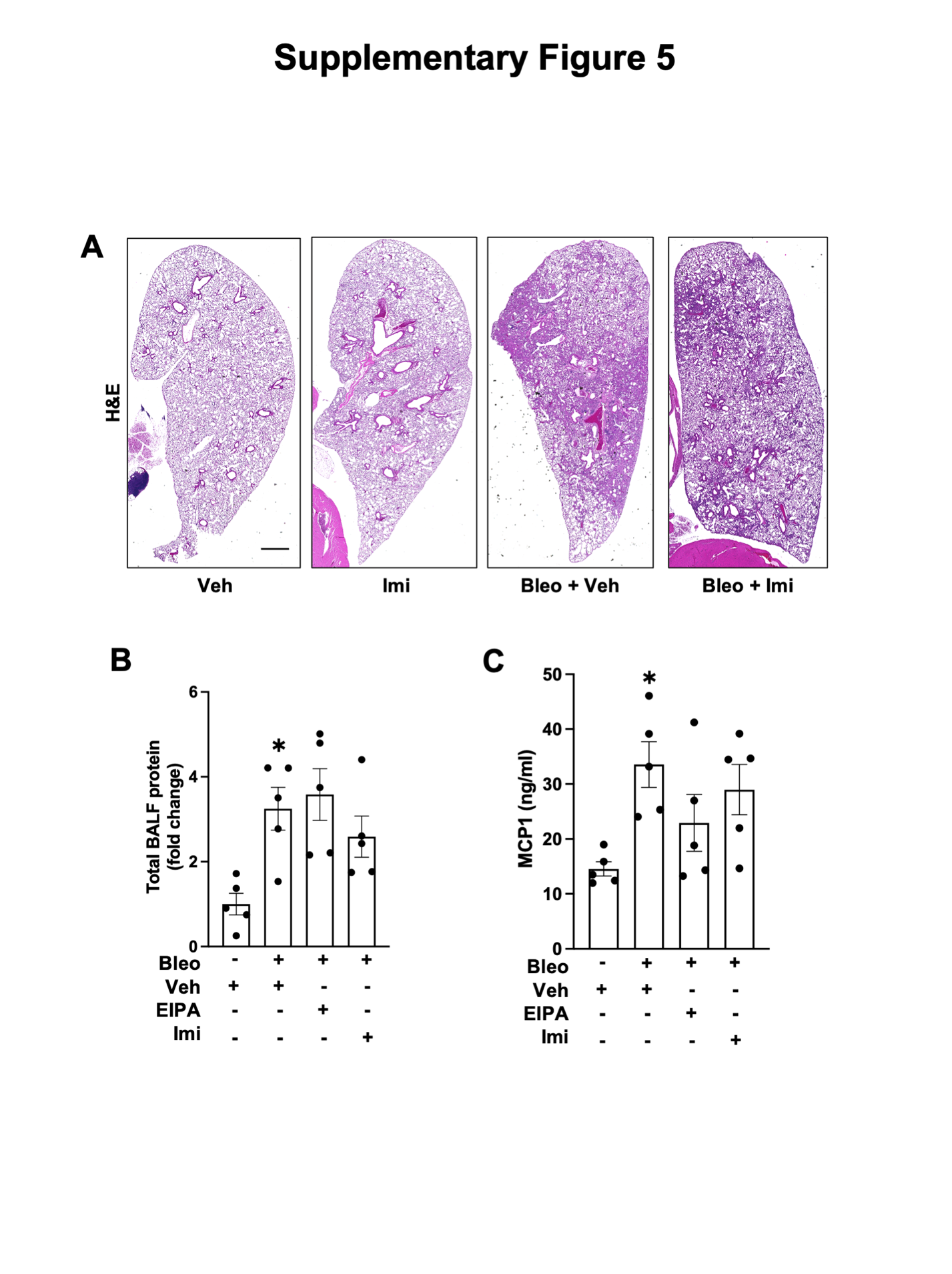

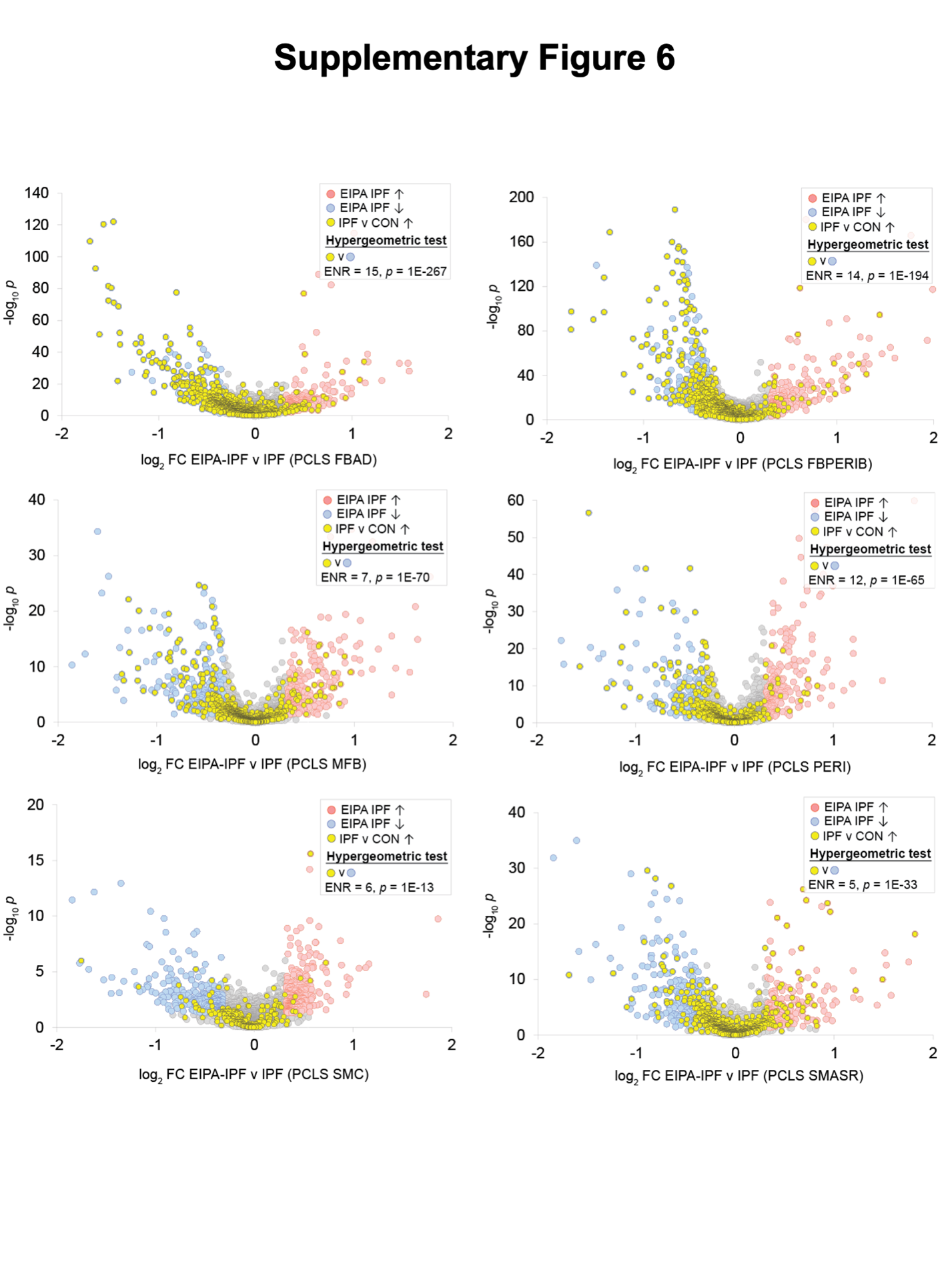

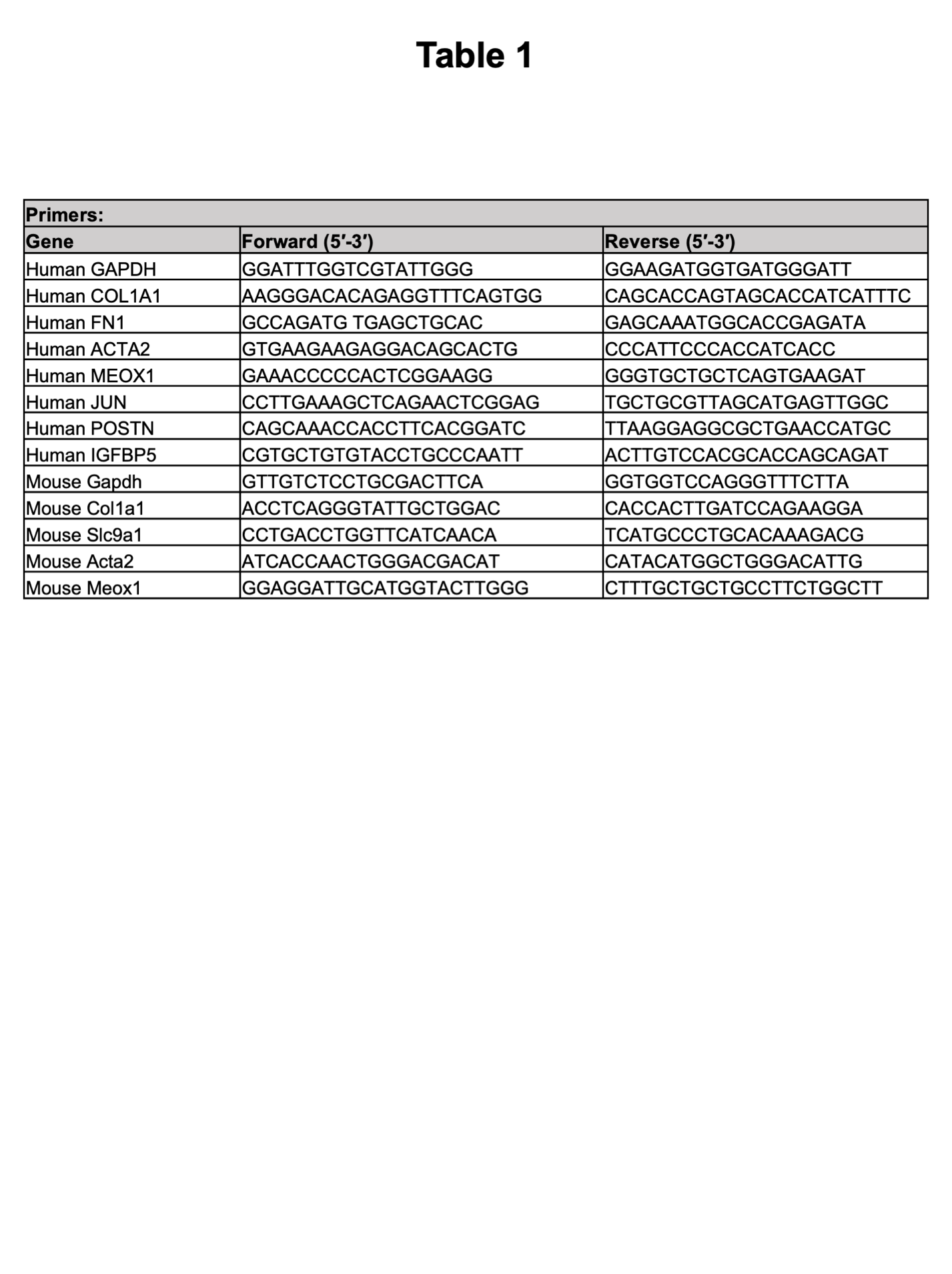
